# Androgen blockade primes NLRP3 in macrophages to induce tumor phagocytosis

**DOI:** 10.1101/2023.09.15.557996

**Authors:** Kiranj Chaudagar, Srikrishnan Rameshbabu, Shenglin Mei, Taghreed Hirz, Ya-Mei Hu, Anna Argulian, Brian Labadie, Kunal Desai, Sam Grimaldo, Doga Kahramangil, Rishikesh Nair, Sabina DSouza, Dylan Zhou, Mingyang Li, Farah Doughan, Raymond Chen, Jordan Shafran, Mayme Loyd, Zheng Xia, David B. Sykes, Amy Moran, Akash Patnaik

**Affiliations:** Section of Hematology/Oncology, Department of Medicine, University of Chicago, Chicago, IL, USA; Center for Regenerative Medicine, Massachusetts General Hospital Cancer Center, Boston, MA, USA; Computational Biology Program, Oregon Health & Science University, Portland, OR, USA; Department of Biomedical Engineering, Oregon Health & Science University, Portland, OR, USA; Department of Molecular Microbiology and Immunology, Oregon Health & Science University, Portland, OR, USA; Department of Cell, Developmental and Cancer Biology, Oregon Health and Science University, Portland, OR, USA; Knight Cancer Institute, Oregon Health and Science University, Portland, OR, USA

## Abstract

Immune-based therapies induce durable remissions in subsets of patients across multiple malignancies. However, there is limited efficacy of immunotherapy in metastatic castrate-resistant prostate cancer (mCRPC), manifested by an enrichment of immunosuppressive (M2) tumor-associated macrophages (TAM) in the tumor immune microenvironment (TME). Therefore, therapeutic strategies to overcome TAM-mediated immunosuppression are critically needed in mCRPC. Here we discovered that NLR family pyrin domain containing 3 (NLRP3), an innate immune sensing protein, is highly expressed in TAM from metastatic PC patients treated with standard-of-care androgen deprivation therapy (ADT). Importantly, *ex vivo* studies revealed that androgen receptor (AR) blockade in TAM upregulates NLRP3 expression, but not inflammasome activity, and concurrent AR blockade/NLRP3 agonist (NLRP3a) treatment promotes cancer cell phagocytosis by immunosuppressive M2 TAM. In contrast, NLRP3a monotherapy was sufficient to enhance phagocytosis of cancer cells in anti-tumor (M1) TAM, which exhibit high *de novo* NLRP3 expression. Critically, combinatorial treatment with ADT/NLRP3a in a murine model of advanced PC resulted in significant tumor control, with tumor clearance in 55% of mice via TAM phagocytosis. Collectively, our results demonstrate NLRP3 as an AR-regulated “macrophage phagocytic checkpoint”, inducibly expressed in TAM by ADT and activated by NLRP3a treatment, the combination resulting in TAM-mediated phagocytosis and tumor control.

## INTRODUCTION

Approximately 1.4 million men are diagnosed with prostate cancer (PC) annually, which accounts for more than 375,000 annual deaths worldwide^1^. While intensified androgen deprivation therapy (ADT) with or without chemotherapy represents the standard-of-care for metastatic prostate cancer (PC), progression to metastatic castration-resistant prostate cancer (mCRPC) is inevitable, causing significant morbidity and mortality^2,3^. Although there has been unprecedented success of immune checkpoint inhibitor (ICI) therapy within subsets of patients across a range of malignancies^4^, efficacy has been limited in mCRPC^5–7^. One of the critical factors limiting ICI efficacy in mCRPC is an enrichment of immunosuppressive tumor-associated macrophages (TAM) within the tumor microenvironment (TME)^8–10^.

The therapeutic activation of Pattern Recognition Receptors, such as cytosolic NOD-like receptors (NLR), modulates the innate immune system to induce proinflammatory cytokine release, including Type I interferons (IFN), resulting in anti-tumor immune responses^11^. NLR family pyrin domain containing 3 (NLRP3) is an intracellular multimolecular protein complex which stimulates cleavage of pro-caspase-1 into active caspase-1. This results in activation of IL- 1β, IL-18, and Gasdermin D, the latter being a pore-forming protein that facilitates cytokine release and pyroptosis, a form of inflammatory cell death^12^. Successful NLRP3 activation requires the presence of two signals. The first is a priming signal, which induces the nuclear factor kappa B (NF-kB) pathway to upregulate both NLRP3 expression and pro-IL1β. The second, an activation signal, can be initiated by cellular stress signals and the sensing of pathogen and damage-associated molecular patterns (PAMP/DAMP)^12^. The consequence of NLRP3 inflammasome activation on tumorigenesis is complex, and implicated in both pro-tumorigenic^13,14^ and anti- cancer processes^11^. Studies in immunogenic murine models showed that NLRP3 activation and cytokine release via dendritic cells can result in CD8^+^T cell-dependent IFNγ production and tumor regression as well as caspase-1-dependent pyroptosis and inhibition of autophagy directly in cancer cells^11,15^. Conversely, NLRP3 activation in tumor cells^13^ or cancer-associated fibroblasts (CAFs)^14^ has been shown to result in the recruitment of MDSCs into the TME, thereby promoting tumorigenesis.

Using single cell transcriptomic analysis of patients with metastatic PC (mPC), we demonstrated that NLRP3 was highly expressed in TAM following treatment with ADT. Based on these findings, we hypothesized that androgen axis blockade could enhance NLRP3 expression and potentiate innate immune tumor control in advanced PC. Critically, we discovered that blockade of TAM-intrinsic androgen receptor (AR) activity enhanced NLRP3 expression, but not inflammasome activity in the immunosuppressive (M2) TAM. In contrast, anti-tumor (M1) TAM exhibited high *de novo* NLRP3 expression, regardless of AR activity. The combination of AR blockade and NLRP3 agonism significantly enhanced phagocytosis of cancer cells by M2 TAM, whereas NLRP3 agonist treatment alone was sufficient to induce phagocytosis in M1 TAM. Following AR blockade/NLRP3 agonist combination treatment, all TAM acquired a distinct phenotype with high PD-L1 and CD86 expression, indicative of phagocytic TAM. Critically, NLRP3 agonism in combination with ADT resulted in significant tumor control in an aggressive c-myc driven advanced PC model, with 55% of mice achieving complete tumor clearance, which was abrogated by concurrent clodronate (phagocytic macrophage depletion) treatment, suggesting TAM phagocytosis as a driver of response. Collectively, our results identify the NLRP3 as an AR-regulated “macrophage phagocytic checkpoint” that can be inducibly expressed and activated in TAM following ADT and NLRP3 agonist treatment, respectively.

## RESULTS

### NLRP3 is highly expressed within TAM of mPC

With the goal of discovering novel therapeutic vulnerabilities in the mPC TME, we investigated NLRP3 expression in patients with metastatic prostate cancer who received ADT as part of standard-of-care therapy. Three tissue compartments were sampled from this patient cohort and analyzed by single cell-RNA sequencing (scRNAseq)^16^: (1) tumor = bone metastatic (BMET) tumor, (2) involved = bone marrow (BM) at the vertebral level adjacent to tumor site and (3) distal = BM from a different vertebral body distant from the tumor site but within the surgical field. There was a significant enrichment of NLRP3 expression and inflammasome activity within BMET PC patients, particularly in the myeloid cells, relative to benign BM controls (Fig. 1a, Extended Data Fig. 1a). Among these myeloid populations, NLRP3 expression and inflammasome activity were highest within tumor-inflammatory monocytes (TIM) and tumor-associated macrophages (TAM) at BMET tumor sites, relative to either involved, distal or benign BM controls (Extended Data Fig. 1b, c). Given the significantly increased NLRP3 expression within the TIM and TAM at mPC BMET tumor site, we compared relative expression within TIM and TAM populations across 6 additional cancer types, including untreated primary localized PC (pPC)^16^. Although there was no significant difference in the TAM/TIM frequency across 7 cancer types (Extended Data Fig. 1d, e), NLRP3 expression was highest within the TIM and TAM of patients with BMET PC as compared to 6 other cancer types, including untreated localized primary PC (pPC, Fig. 1b, Extended Data Fig. 1f). Notably, the inflammasome activity of TIM and TAM was similar in BMET PC and in pPC (Fig. 1b, Extended Data Fig. 1g), indicating that enhanced NLRP3 expression alone is insufficient to drive inflammasome activation. Collectively, these data demonstrate that NLRP3 is differentially expressed in macrophage/monocytic lineage of mPC patients, but not in other cancers or cell types within the BMET PC microenvironment.

**Fig. 1.**
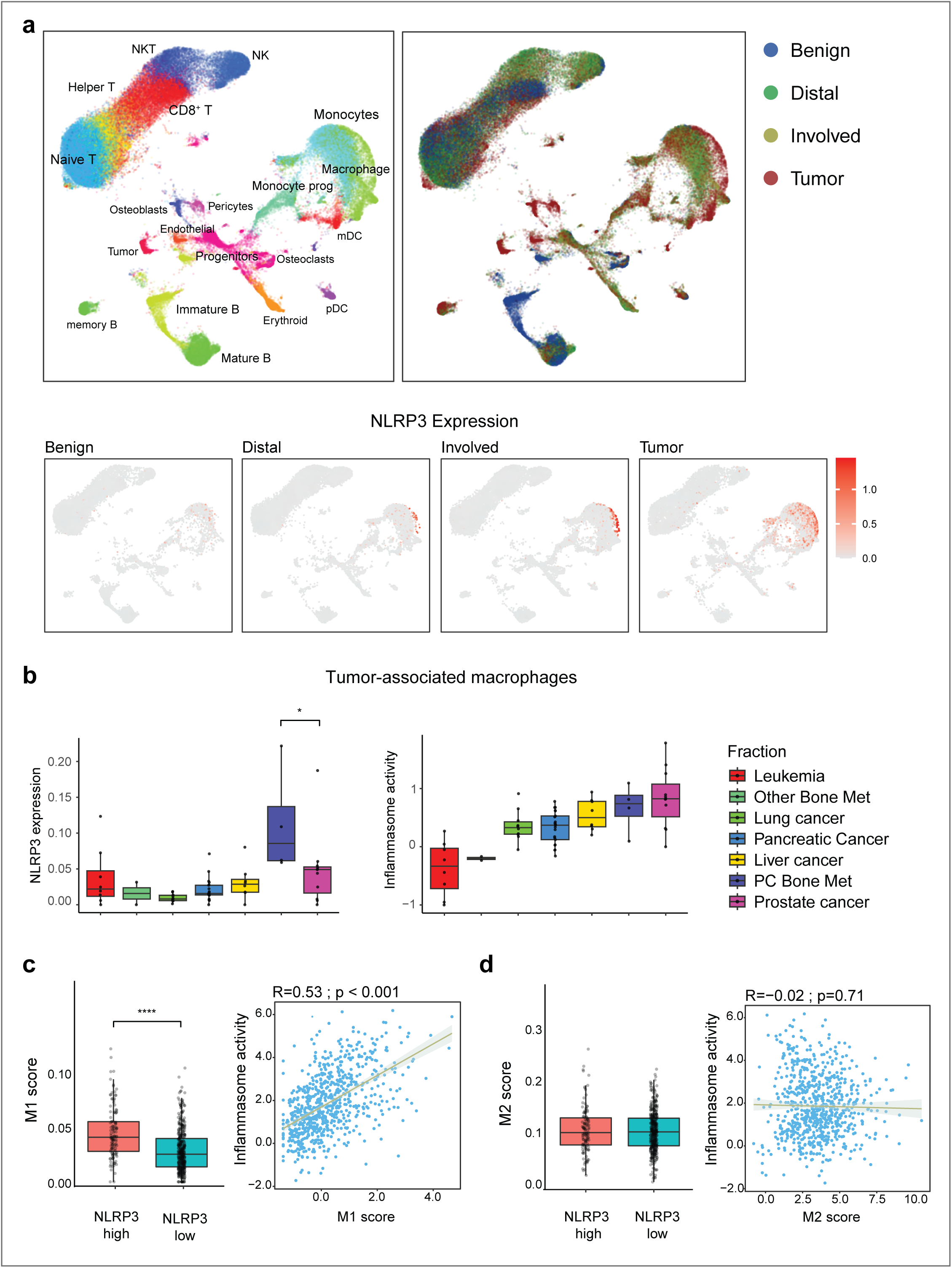
NLRP3 expression is enhanced in tumor-associated macrophages (TAM) of ADT- treated metastatic PC patients, relative to other cancers and untreated primary PC. **a,** Single cell RNAseq (scRNAseq) was performed on biopsies collected from tumor, involved and distant sites of bone metastatic prostate cancer (BMET PC) patients as described in the Methods section, and compared with healthy benign bone marrow. UMAP plots represent cell subpopulations (top left panel) across different anatomic locations of biopsy samples (top right panel). NLRP3 expression was analyzed and visualized in UMAP plots (bottom panel). **b,** scRNAseq was performed to generate boxplots depicting NLRP3 expression and inflammasome activity within TAM across 7 different cancer types listed in the panel. c-d, BMET PC TAM were stratified into two groups based on TAM NLRP3 expression (NLRP3-high: top 25% versus NLRP3-low: bottom 25%), scored for M1 (c)/M2 (d) phenotypes and correlated with inflammasome activity. Data obtained from specimens of BMET PC (n = 7 patients) for panels a-d, healthy benign bone marrow (n = 7 human donors) for panel a, leukemia (n = 20 patients), other bone met (non-prostate bone metastatic cancers, n = 3 patients), lung cancer (n = 10 patients), pancreatic cancer (n = 21 patients), liver cancer (n = 8 patients) and primary prostate cancer patients (n = 11 patients) for panel b. Except panel a, all panels show biological replicates as mean ± s.e. Significances/p-values were calculated by Wilcoxon rank-sum test and indicates as follows, *p<0.05 and ****p<0.0001.

Given that enhanced NLRP3 expression in mPC did not correlate with increased inflammasome activation, we tested the hypothesis that the NLRP3 agonist (NLRP3a) could enhance inflammasome pathway activation in TIM and TAM. We treated immortalized bone marrow derived macrophages (iBMDM, M0-polarized^18^) *in vitro* with NLRP3a (Extended Data Fig. 2a), and observed a dose-dependent increase in markers of inflammasome formation, including cleaved caspase-1 expression and IL-1β/IL-18 secretion, which peaked at approximately 10 μM (Extended Data Fig. 2b, c, d). These data demonstrate that the NLRP3 small molecule agonist can drive NLRP3 pathway activation in macrophages *ex vivo*.

### M1 TAM exhibit high NLRP3 expression, but not M2 TAM

TAMs are heterogenous with respect to their activation and functionality^19^ and can be broadly classified as M1 (anti-tumor) and M2 (immunosuppressive). To analyze NLRP3 expression between M1 and M2 TAM, we performed flow cytometry and western blot analysis, which revealed higher expression of NLRP3 and immunostimulatory markers, MHC-II and CD86^20^, and lower expression of immunosuppressive markers, Arg1 and CD206^20^ in M1-polarized iBMDM, relative to M2-polarized iBMDM (Extended Data Fig. 3a, b). As expected, these M1 iBMDM also had higher expressions of exhaustion markers PD-1 and PD-L1^20^, and phagocytosis- stimulatory checkpoints, SIRPα, CD16 and CD64^20^, relative to M2 iBMDM (Extended Data Fig. 3a, b). Using single cell transcriptomic analysis in BMET PC, we observed that TAM with high NLRP3 expression also had a higher M1 score. Furthermore, there was a positive correlation between M1-score and inflammasome activity (Fig. 1c), not observed in M2 TAM (Fig. 1d).

As the first step to elucidate the impact of acute NLRP3a treatment on the expression of NLRP3 across different immune cell lineages *in vivo*, we utilized the murine syngeneic Myc-CAP model of PC, which recapitulates the molecular and immunological landscape of advanced PC and is also known to be refractory to ICI^21^. To compare our human patient data to our Myc-CAP murine model, we performed flow cytometry to identify NLRP3-expressing populations in the TME (Extended Data Fig. 4a). Consistent with the human PC BMET data (Fig. 1a, b, Extended Data Fig. 1b), we found higher NLRP3 expression within TAM at both early (17-fold) and late (12- fold) phases of tumor growth, when compared to the CD45^-^ fraction. In addition, Gr-MDSC (13- and 7.5-fold at early and late phases, respectively) and CD4^+^T cells (10- and 8-fold at early and late phases, respectively) had high NLRP3 expression at both phases of tumor growth while cancer-associated fibroblasts (CAF, 18-fold) demonstrated high NLRP3 expression only at early phase (Extended Data Fig. 4b, c). Next, Myc-CAP syngeneic tumors were harvested at 24, 48, and 72 hours following a single intra-tumoral (*it*) dose of NLRP3a (Extended Data Fig. 5a). Despite high intrinsic baseline NLRP3 expression in CD4^+^ T cells, Gr-MDSCs, and TAM in untreated tumors, M1 TAM (MHC-II^+^ TAM, Extended Data Fig. 3a) sustained higher NLRP3 expression relative to Gr-MDSCs, CD4^+^ T cells and M2 TAM following 72 hours of acute NLRP3a treatment (Extended Data Fig. 5b, c). While the frequency of M1 TAM and CD4^+^ T cells did not decrease following NLRP3 activation within TME, there was an approximately 15-fold and 3-fold increase in Gr-MDSC and M2 TAM infiltration, respectively, following 72 hours of acute NLRP3a treatment (Extended Data Fig. 5d, e). Collectively, these data demonstrate that M1 TAM exhibit high NLRP3 expression which is sustained following acute NLRP3 agonist treatment *in vivo*, and accompanied by an increase in NLRP3-low Gr-MDSC and M2-TAM.

### AR inhibition primes NLRP3 within M2 TAM

We observed enhanced NLRP3 expression within the TAM of mPC, relative to other tumor types and primary PC (Fig. 1b). Since all mPC patients had been treated with ADT (gonadotropin-releasing hormone [GnRH] agonist with or without androgen receptor signaling inhibitor) as compared to patients with primary PC and non-prostate malignancies who did not receive ADT (Fig. 1b), we hypothesized that AR axis blockade might enhance TAM NLRP3 expression. Using bulk-RNAseq dataset of patients treated with AR inhibitor (ARi) enzalutamide for metastatic PC, we observed an inverse relationship of AR activity with NLRP3 expression (Fig. 2a) and with inflammasome activity (Fig. 2b), which was preserved across lymph node, bone, and liver metastases. Furthermore, we observed a direct correlation between response to pembrolizumab following enzalutamide progression and high M1 macrophage activity within the TME, relative to the non-responder subset (Fig. 2c). We performed a similar analysis of the West Coast Dream Team (WCDT) mCRPC cohort (n=99) who had been treated with various androgen axis inhibitors and noted a consistent inverse correlation of AR activity with inflammasome activity (Fig. 2d). Consistent with these human bioinformatic analyses, our *ex vivo* murine studies demonstrated high AR/FKBP5 expression with corresponding low NLRP3 level in M2-iBMDM, relative to M1- iBMDM (Extended Data Fig. 3b). Collectively, these findings demonstrate an inverse relationship between NLRP3 expression and AR activity, with high NLRP3 expression associated with an M1 signature and a favorable clinical response to ICI in mCRPC.

**Fig. 2.**
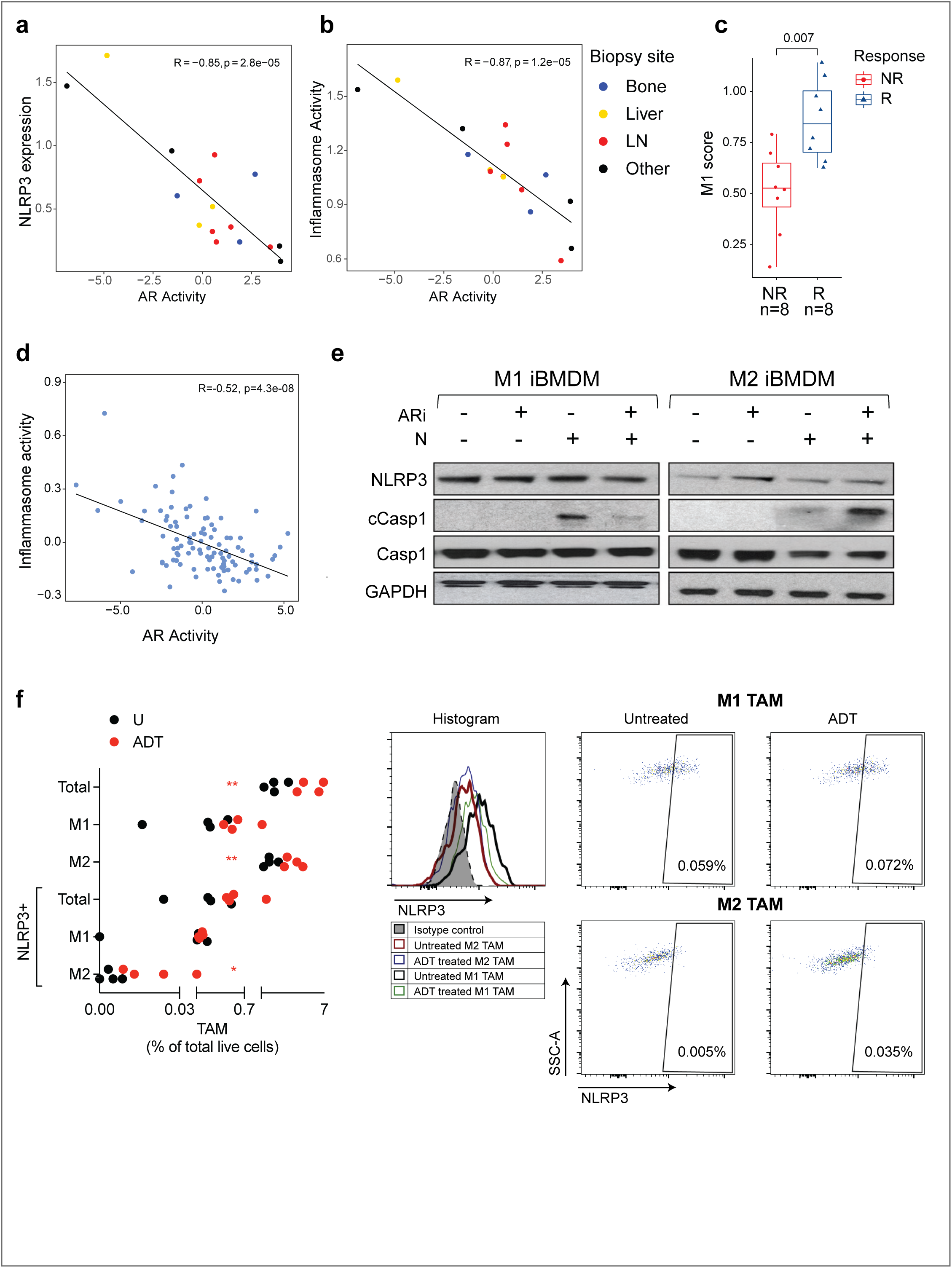
AR blockade primes NLRP3 in immunosuppressive M2 macrophages, but not in anti-tumor M1 macrophages. **a-c**, Patients with mCRPC who had previously progressed on enzalutamide were treated with pembrolizumab (anti-PD1 antibody) in the context of an investigator-initiated clinical trial (NCT02312557). Bulk RNAseq data on baseline metastatic biopsy samples from enzalutamide-refractory mCRPC patients prior to pembrolizumab treatment were utilized to correlate AR activity with NLRP3 expression (**a**)/inflammasome activity (**b**) and represented using scatter plots. The mCRPC patients were stratified into responder (R) *vs* non- responder (NR) groups based on PSA response as described in the Methods section, and the M1 score estimated using bulk RNAseq (**c**). **d**, Correlation analysis of AR activity with NLRP3 expression was performed using bulk RNAseq dataset available from West Coast Dream Team (WCDT) data repository. **e**, iBMDM were differentiated into M1 and M2 phenotypes and treated with enzalutamide (ARi, 10μM, 24 hours), NLRP3a (10μM, 1 hour) or their combination. The indicated markers of inflammasome activation were probed by western blotting. **f**, Syngeneic FVB mice with established Myc-CAP tumors (∼200 mm^3^) were treated with single dose of degarelix (ADT, 0.625μ*g*, *sc*) for 7 days. Size-matched untreated and treated tumors were profiled to assess total and NLRP3-expressing TAM frequency within the TME. Representative flow cytometry plots depict TAM-NLRP3 expression and frequency of NLRP3^+^TAM (relative to total live cells within TME). Data obtained from metastatic PC (n = 16 patients) for panel **a**, **b**, **c**, metastatic PC (n = 111 patients) for panel **d**, n = 3 biological replicates for *in vitro* experiments in panel **e** and n = 4 mice per group for *in vivo* experiments in panel **f**. With the exception of panel **e**, all panels show biological replicates as mean ± s.e. Significances/p-values were calculated by one-way ANOVA test (panel **f**), Kruskal-Wallis test (panel **a**, **b**, **c**, **d**) and indicates as follows, *p<0.05 and **p<0.01.

Since we observed a direct correlation between AR axis blockade and NLRP3 expression, we hypothesized that AR signaling may suppress NLRP3 expression within TAM. To assess the impact of AR blockade on NLRP3 expression and inflammasome activation within TAM *in vitro*, we performed western blotting on M1 and M2 differentiated iBMDM following treatment with the testosterone analogue R1881 and the AR inhibitor enzalutamide. Consistent with the *in vivo* findings, NLRP3 expression increased following enzalutamide treatment in M2 iBMDM, which translated to enhanced inflammasome pathway activation (cleaved caspase-1 expression) in combination with NLRP3a (Fig. 2e). Conversely, there was no increase in NLRP3 expression in the enzalutamide-treated M1 iBMDM, and a paradoxical decrease in cleaved caspase-1 expression with the combination of NLRP3a and enzalutamide (Fig. 2e).

Next we profiled NLRP3 expressing TAM in Myc-CAP following ADT (degarelix, a GnRH antagonist), and observed an increase in TAM infiltration, particularly M2 TAM within the TME at peri-castration (day 7, ∼200 mm^3^) stage (untreated tumor size of ∼200 mm^3^, Fig. 2f). Although there was a lower frequency of M1 TAM (∼0.22%) relative to M2 TAM (∼1.58%) within the TME at baseline, we observed higher NLRP3 expressing M1 TAM (∼0.07%) relative to M2 TAM (∼0.004%, Fig. 2f). Degarelix treatment resulted in a significant increase in NLRP3- expressing M2 TAM (∼0.01%), but not in M1 TAM (∼0.09%, Fig. 2f), consistent with *de novo* high NLRP3 expression within M1 TAM and ADT-induced increase in M2 TAM within BMET PC (Fig. 1c, d). Collectively, these data demonstrate that AR blockade can prime NLRP3 expression within immunosuppressive M2 TAM *in vitro* and *in vivo*, thus creating a therapeutic vulnerability for NLRP3 activation in this immunosuppressive subset (in addition to anti-tumor M1 TAM) within the TME. (Figure 2f).

### AR inhibitor/NLRP3 agonist treatment drives iBMDM-mediated cancer cell phagocytosis

To elucidate the phenotypic and functional consequences of AR-blockade induced NLRP3 priming in immunosuppressive M2 macrophages, we performed repolarization and phagocytosis assays respectively. M1 and M2 iBMDM were treated with enzalutamide alone and in combination with NLRP3 agonist. At baseline, we observed a higher expression of the suppressive markers, Arg1 and CD206, and a lower expression of activation markers, MHC-II and CD86 within M2 iBMDM, relative to M1 iBMDM (Fig. 3a, b, Extended Data Fig. 3a, 6a, b, c). Enzalutamide suppressed both Arg1 and CD206 expression on M2 iBMDM, which was accentuated by concurrent treatment with NLRP3a for 1 hour, particularly for CD206 (Fig. 3a, b). However, neither enzalutamide, NLRP3a nor their combination increased MHC-II and CD86 on M2 iBMDM (Extended Data Fig. 6a, b, c), demonstrating that enzalutamide/NLRP3a combination specifically inhibits the immunosuppressive phenotype of M2 iBMDM, but does not polarize them towards M1 TAM. Treatment with either single agents or their combination did not alter Arg1, CD206, MHC-II and CD86 expression on M1 iBMDM (Fig. 3a, b, Extended Data Fig. 6a, b, c). Importantly, NLRP3a treatment for 1-hour or for 24-hours resulted in an increase in phagocytosis of cancer cells within M1 iBMDM, which was not accentuated by concurrent enzalutamide treatment (Fig. 4a, b). In contrast, NLRP3a monotherapy (1-hour), or in combination with enzalutamide, did not enhance phagocytosis of Myc-CAP tumor cells by M2 iBMDM. However, after 24 hours of NLRP3a treatment, we observed enhanced phagocytosis of cancer cells within M2 iBMDM, which was significantly enhanced by enzalutamide, and equivalent to enzalutamide/NLRP3a treated M1 iBMDM (Fig. 4a, b). Collectively, these data demonstrate that AR blockade-mediated NLRP3 priming synergizes with NLRP3a treatment to enhance phagocytosis of cancer cells by M2 macrophages. Since anti-tumor M1 macrophages are already primed for high NLRP3 expression, enhancement of phagocytosis is achieved with NLRP3 monotherapy alone, obviating the need for ADT-induced NLRP3 priming.

**Fig. 3.**
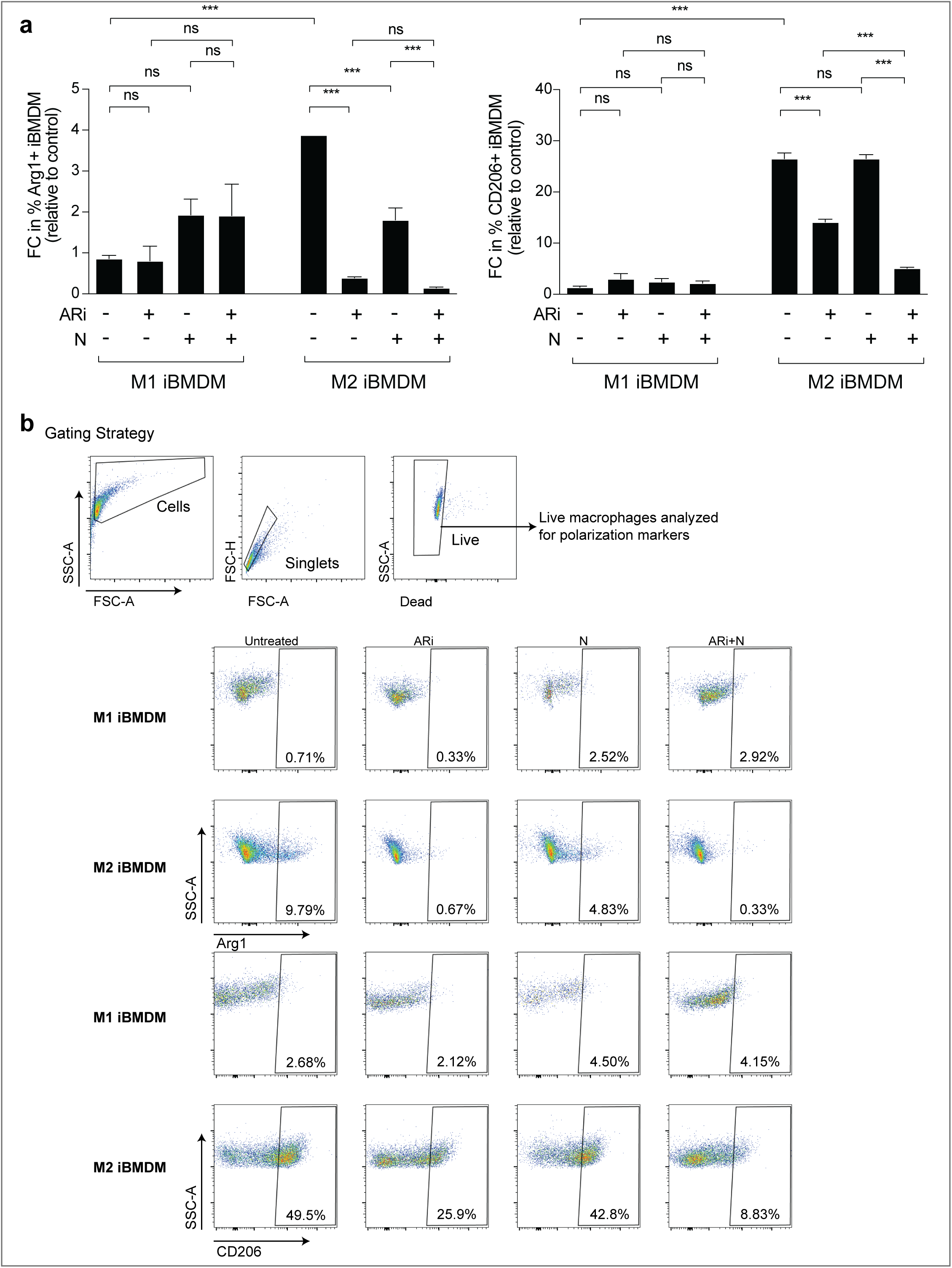
*In vitro* AR blockade/NLRP3 agonist treatment overcomes suppressive status of M2 iBMDM. **a-b**, iBMDM were differentiated into M1 and M2 phenotypes using cytokine cocktails, as described in the Methods section and treated with enzalutamide (ARi, 10μM, 24 hours), NLRP3a (10μM, 1 hour), or their combination. Flow cytometry was performed to identify Arg1 (Left panel) and CD206 (Right panel) expressing iBMDM (**a**). Representative flow cytometry plots depict gating strategy of live M1/M2 iBMDM and quantification of their suppressive status following *in vitro* ARi+/-NLRP3a treatment vs untreated control, defined as Arg1 or CD206 expressing iBMDM relative to total live M1/M2 iBMDM (**b**). Data obtained from n=3 biological replicates. Panel **a** shows biological replicates as mean ± s.e. Significances/p-values were calculated by one-way ANOVA and indicates as follows, ***p<0.001; ns = not statistically significant. FC = fold change. Control = untreated M1 iBMDM group.

**Fig. 4.**
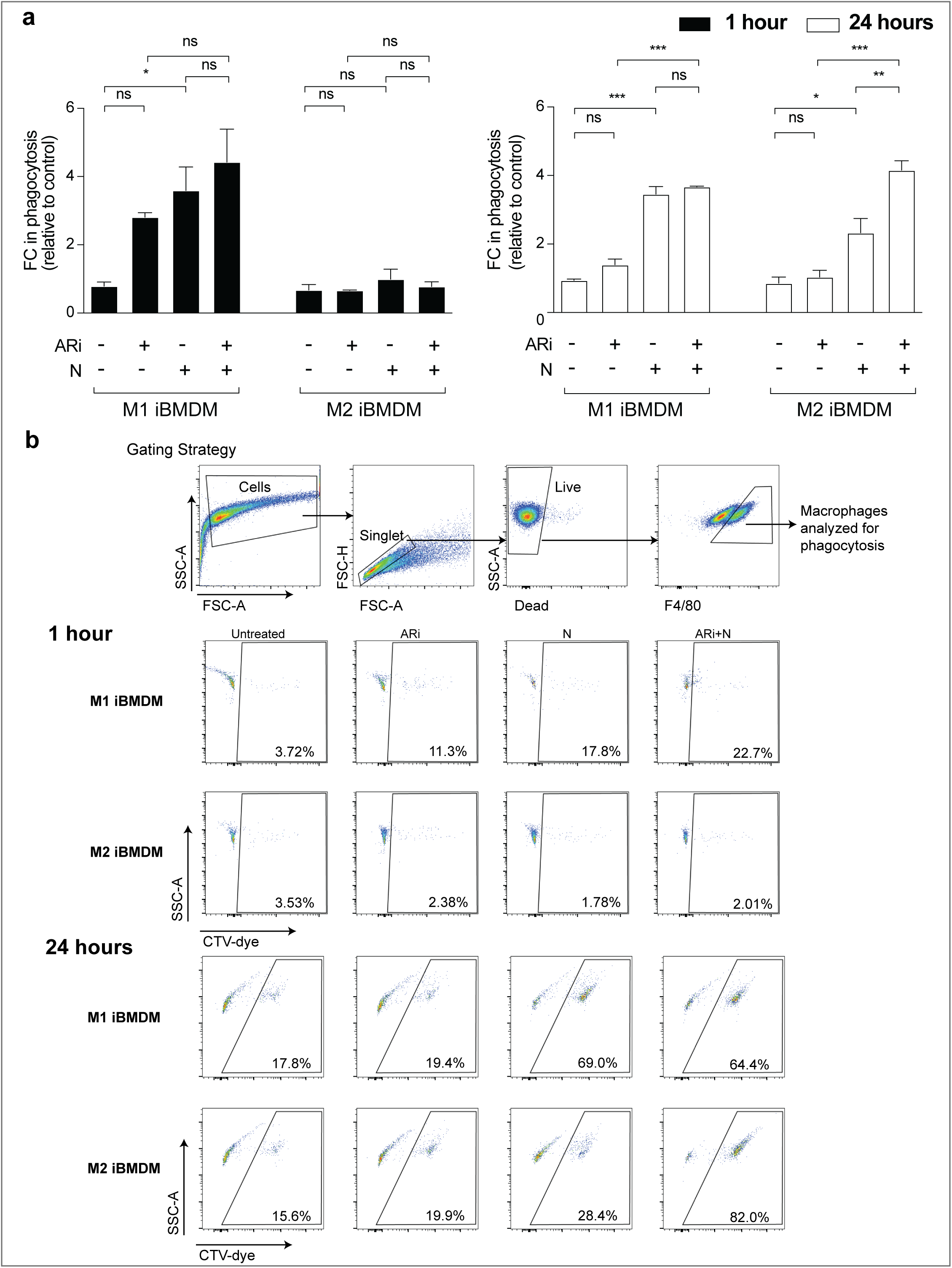
*In vitro* AR blockade/NLRP3 agonist treatment M1/M2 iBMDM induces phagocytic uptake of Myc-CAP tumor cells by NLRP3 primed phagocytic (NPP) iBMDM. **a-b,** iBMDM were differentiated into M1 and M2 phenotypes using cytokine cocktail, as described in the Methods section, and treated with enzalutamide (ARi, 10μM, 24 hours), NLRP3a (10μM, 1 hour or 24 hours), or their combination following co-culture with CTV-dye stained Myc-CAP cells for 1 hour. Flow cytometry was performed to determine CTV-dye^+^iBMDM, indicative of phagocytosis of tumor cells, and their phagocytic status was determined as a percentage of total live M1/M2 iBMDM. These data were normalized to untreated N1 iBMDM group to calculate fold change in phagocytosis (**a**). Representative flow cytometry plots depict gating strategy of live M1/M2 iBMDM and their phagocytic status (as a percentage of total live M1/M2 iBMDM) following *in vitro* ARi+/-NLRP3a treatment (**b**). Data obtained from n=3 biological replicates. Panel **a** shows biological replicates as mean ± s.e. Significances/p-values were calculated by one-way ANOVA and indicates as follows, *p<0.05, **p<0.01 and ***p<0.001; ns = not statistically significant. FC = fold change. Control = untreated M1 iBMDM group.

As both M1 and M2 macrophages demonstrate enhanced phagocytosis following treatment with ADT and NLRP3 agonist and there is no subsequent change in polarization, we next performed flow cytometric profiling with 11 different TAM phenotypic/functional markers to characterize this macrophage subset, so it could be investigated *in vivo* following NLRP3 agonism based therapy in the murine Myc-CAP model. Following NLRP3a/enzalutamide combination treatment, we observed no change in PD-L1 and CD86 marker upregulation or phagocytosis at 1 hour (Fig. 4a, b, 5a). However, there was a significant increase in PD-L1 and CD86 expression on NLRP3-primed phagocytic (NPP) iBMDM derived from either M1 or M2 iBMDM following 24-hour treatment with NLRP3a/enzalutamide combination, which was associated with enhanced phagocytosis (Fig. 5a, b, c). Consistent with our data in Fig. 4b, NLRP3 monotherapy increased PD-L1 and CD86 marker expression in M1 iBMDM at 24 hours, which was not enhanced further with concurrent enzalutamide therapy (Fig. 5b, c). Importantly, we observed no change in PD-L1 expression following single-agent treatment with enzalutamide or NLRP3 agonist or their combination in the absence of tumor cell co-culture with total M1/M2 iBMDM (Extended Data Fig. 7). However, M1 iBMDM even in absence of co-culture exhibited enhanced CD86 expression (consistent with early emergence of macrophage activation) following NLRP3a monotherapy, which was not accentuated with combinatorial treatment (Extended Data Fig. 7). Critically, we observed no change in PD-L1 and CD86 expression in the M1/M2 non-phagocytic (np) iBMDM (Fig. 5d, e). Taken together, these data validate PD-L1 and CD86 as phenotypic biomarkers of NPP iBMDM cells that exhibit enhanced phagocytosis of tumor cells.

**Fig. 5.**
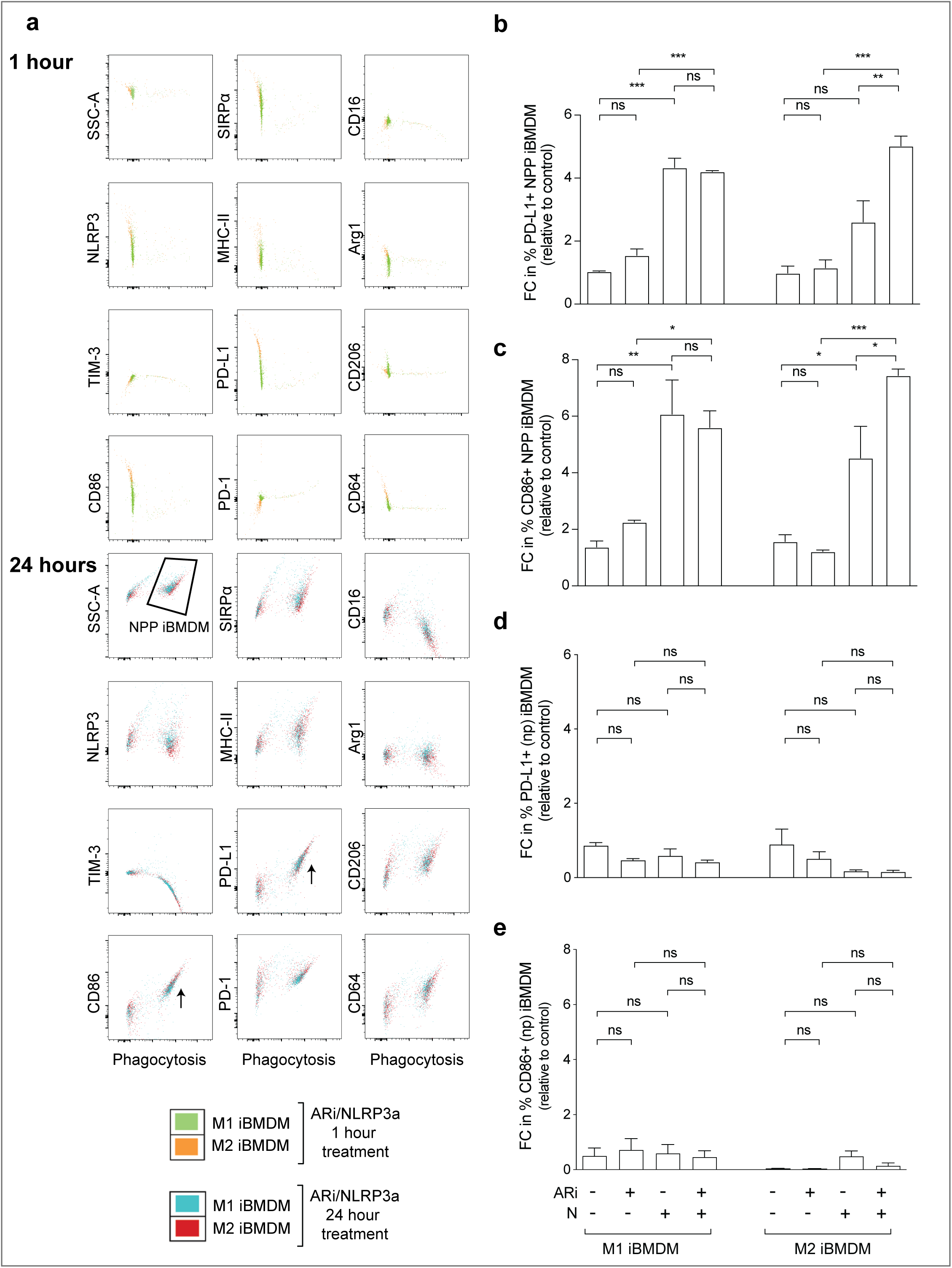
NLRP3 primed phagocytic (NPP) iBMDM express PD-L1 and CD86. **a-e**, iBMDM were differentiated into M1 or M2 phenotypes using cytokine cocktail, as described in the Methods section, and treated with enzalutamide (ARi, 10μM, 24 hours), NLRP3a (10μM, 1 hour or 24 hours) or their combination, and co-cultured with CTV-dye stained Myc-CAP cells for 1 hour. Flow cytometry was performed to identify markers associated with NPP iBMDM-mediated phagocytic uptake. Representative flow cytometry plots depicting surface or intracellular marker expression on Y-axis and CTV-dye/phagocytic uptake on X-axis. If markers are either positively or negatively associated with phagocytosis, NPP iBMDM (right side cluster on X-axis) shifts up or down, respectively, relative to non-phagocytic (np) M1/M2 iBMDM (left side cluster on X-axis, (**a**)). NPP iBMDM and (np) iBMDM were quantified for PD-L1 (**b** and **d**, respectively) and CD86 (**c** and **e**, respectively) expression status. Data obtained from n=3 biological replicates. Panel **b-e** show biological replicates as mean ± s.e. Significances/p-values were calculated by one-way ANOVA and indicates as follows, *p<0.05, **p<0.01 and ***p<0.001; ns = not statistically significant. FC = fold change. Control = untreated M1 iBMDM group. Kaplan-Meier survival analysis (panel **b**) and indicates as follows, *p<0.05, **p<0.01 and ***p<0.001; ns = not statistically significant. U = untreated tumors.

### NPP TAM eradicates cancer *in vivo*

Based on the observed induction of tumor cell phagocytosis *ex vivo* by ARi/NLRP3a- induced NPP iBMDM, we hypothesized that this combinatorial approach would be therapeutically beneficial in the syngeneic Myc-CAP murine model. Flow cytometric analysis of the untreated Myc-CAP TME revealed 2.1% of TAM, approximately 1/5 of which are PD-L1^+^ or CD86+ NPP- TAM (Fig. 6a), which are insufficient to control tumor growth.

**Fig. 6.**
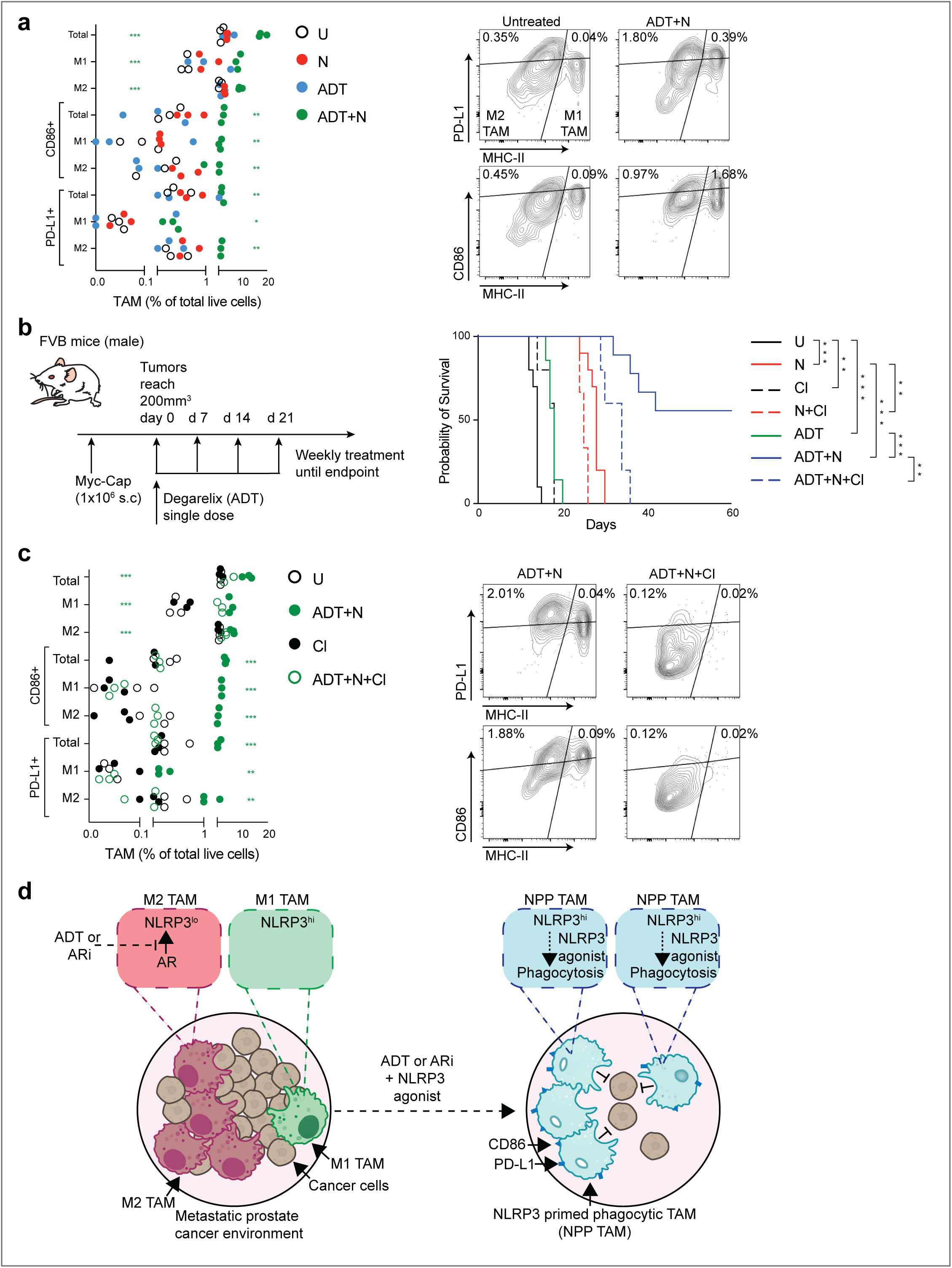
ADT/NLRP3 agonist-induced NLRP3 primed phagocytic-tumor associated macrophages (NPP-TAM) drive tumor clearance in the majority of Myc-CAP tumor-bearing syngeneic mice. **a**, Syngeneic FVB mice with established Myc-CAP tumors (∼200 mm^3^) were treated with single dose of degarelix (ADT, 0.625μ*g*, *sc*), NLRP3a (0.3 mg/kg, *it*) or their combination for 7 days. Harvested tumors were analyzed by flow cytometry to calculate frequency of PD-L1/CD86 expressing TAM within TME (Left panel). Representative flow cytometry plots depicting PD-L1/CD86-expressing NPP TAM (relative to total live cells) within TME following *in vivo* ADT+N treatment vs untreated control (Right panel). **b-c**, Myc-CAP tumor-bearing mice (∼200 mm^3^) were treated with indicated agent(s) either until non-palpable tumors for 4 weeks or until tumors reached euthanasia end-point (∼2500 mm^3^). In parallel, corresponding cohorts of mice received clodronate (Cl, 200μg, *ip*, 1-hour pretreatment), to systemically deplete phagocytic TAM, prior to administration of N and ADT/N. Survival curves were plotted for 60 days (**b**). Harvested tumors were analyzed by flow cytometry to quantify PD-L1/CD86 expressing TAM within TME (Left panel, (**c**)). Representative flow cytometry plots depicting PD-L1/CD86-expressing NPP TAM (relative to total live cells) within TME following *in vivo* ADT+N+Cl treatment as compared to ADT+N group (Right panel, (**c**)). **d**, Schema represents primary or metastatic PC TME which have an enrichment of immunosuppressive M2 TAM and a relative paucity of anti-tumor NLRP3- expressing M1 TAM. AR blockade (ARB) primes NLRP3 in the M2 TAM, and NLRP3a reprograms these NLRP3 primed M1/M2 TAM into PD-L1/CD86 expressing NPP TAM, which phagocytose cancer cells, resulting in tumor clearance. Data obtained from n = 3 mice per group for panel **a** and **c**, and n = 6-9 mice per group for panel **b**. Except panel **d**, all panels show biological replicates as mean ± s.e. Significances/p-values were calculated by two-way ANOVA (panel **a**, **c**),

NLRP3a monotherapy for 1 week increased the population of NPP TAM (∼1.5- and ∼2.1- fold with PD-L1 and CD86 expression, respectively) within the TME, while degarelix alone for 1 week did not (∼0.8- and ∼0.3-fold, respectively) (Fig. 6a). Consistent with our *in vitro* experiments (Fig. 5a, b, c), degarelix/NLRP3a combination therapy for 1 week significantly enhanced NPP TAM (PD-L1 and CD86 expressing cells by ∼3.3- and ∼3.8-fold, respectively) relative to NLRP3a treated group (Fig. 6a), thus demonstrating a synergistic increase in NPP TAM accumulation with combinatorial degarelix and NLRP3a treatment within the TME. Mice treated with NLRP3a monotherapy demonstrated significant tumor control and improved median survival rate (29 days vs. 14 days in control mice, Fig. 6b and Extended Data Fig. 8). The combination of degarelix and NLRP3a led to long-term tumor control (>60 days), with 55% of mice achieving complete tumor clearance (Fig. 6b and Extended Data Fig. 8). Flow cytometric analysis of tumors at endpoint revealed a ∼5.6-fold enrichment of NPP TAM in degarelix/NLRP3a treated group relative to untreated (Fig. 6c), highlighting the potential role of NPP TAM in driving the *in vivo* anti-tumor response. To address the possibility that other immune cell types within the TME may be driving the anti-tumor response following degarelix/NLRP3 treatment, we concurrently depleted NPP TAM using liposomal clodronate (which selectively depletes phagocytic TAM) and observed significant reversal of benefit of degarelix/NLRP3a combination, decreasing the median survival from >60 days to ∼34 days (Fig. 6b). Notably, no mice exhibited a complete response with the triplet therapy of degarelix/NLRP3a/clodronate (Fig. 6b and Extended Data Fig. 8), thus implicating that NPP TAMs are responsible for the ADT/NLRP3 agonist-induced tumor control *in vivo*.

NLRP3 activation results in the formation of the pore-forming protein Gasdermin D, release of IL-1β, IL-18, and pyroptosis, a form of inflammatory cell death^22^. To determine whether the anti-tumor response observed with ADT/NLRP3a is mediated via NLRP3 inflammasome-mediated cytokine release, we used a protein array to quantify cytokines from enzalutamide/NLRP3a-treated M1/M2 iBMDM. At baseline, M1 iBMDM secreted IL-1β, a NLRP3-regulated cytokine^12^ and other pro-inflammatory cytokines (IFNα/β/γ, TNFα, GM-CSF, CXCL1, CXCL10, CCL5, IL-6, IL-12p70), which was not observed in M2 iBMDM (Extended Data Fig. 9a), consistent with our findings that M2 TAM lack inflammasome activity. Treatment of M1 iBMDM with NLRP3a monotherapy resulted in an increase in IL-1β secretion, which was suppressed in combination with enzalutamide (Extended Data Fig. 9b). These findings are consistent with the observed decrease in inflammasome activity in M1 iBMDM with enzalutamide/NLRP3a combination treatment, relative to NLRP3a monotherapy (Fig. 2e). While NLRP3a/enzalutamide combination therapy increased inflammasome activity in M2 iBMDM (Fig. 2e), this was not accompanied by an increase in IL-1β secretion (Extended Data Fig. 9b). Additionally, enzalutamide decreased secretion of IFNβ and IL-10 from M1 iBMDM, which was partially reversed by NLRP3a addition (Extended Data Fig. 9c, d). Enzalutamide alone or in combination with NLRP3a did not alter secretion of these cytokines from M2 iBMDM (Extended Data Fig. 9c, d). Critically, enzalutamide, NLRP3a or their combination did not alter viability or induce pyroptosis^23^ of M1/M2 iBMDM, relative to untreated controls (Extended Data Fig. 10a, b). Collectively, these findings highlight an absence of contribution of inflammasome mediated cytokine release or pyroptosis to the anti-tumor response driven by NPP TAM *in vivo*.

To assess the possible contribution of tumor cell intrinsic NLRP3 activation on tumor control, we looked for NLRP3 expression and activation within cancer cells, following treatment with enzalutamide, singly and in combination with NLRP3 agonist. Myc-CAP cells lacked NLRP3 expression and activation at baseline (Extended Data Fig. 11a, left panel), which was unchanged following enzalutamide, NLRP3a monotherapy or their combination treatment (Extended Data Fig. 11a, right panel). Furthermore, *in vitro* enzalutamide/NLRP3a combination treatment caused a modest decrease in Myc-CAP proliferation (Extended Data Fig. 11b). Critically, pre-treatment of Myc-CAP cells with enzalutamide, NLRP3a monotherapy or their combination treatment did not alter their phagocytosis within M1 or M2 iBMDM (Extended Data Fig. 11c, d), demonstrating that an ADT/NLRP3a-induced tumor cell extrinsic phagocytic mechanism is responsible for tumor control. Taken together, these data demonstrate that androgens directly suppress NLRP3 expression and downstream activation within immunosuppressive M2 TAM, and its abrogation by AR blockade results in significant NPP TAM-mediated tumor control in combination with NLRP3 agonist treatment (Fig. 6d).

## DISCUSSION

Macrophages play an important anti-cancer role via phagocytosis to mount both an innate immune response and to stimulate adaptive immunity through presentation of tumor-specific antigens^24^. Various therapeutic strategies have been developed to induce macrophage phagocytosis, either by inhibition of immunosuppressive TAMs within the TME (e.g., CSF-1 inhibition) or blockade of inhibitory phagocytic checkpoints on tumor cells^25,26^. The latter strategy has demonstrated limited clinical activity^27,28^ and includes agents that target “don’t-eat-me” macrophage inhibitory checkpoints (i.e., CD47, Siglec G and PD-L1) overexpressed by tumor cells to evade phagocytosis^29^. Here we discovered NLRP3 as a novel androgen regulated macrophage phagocytic checkpoint in PC. Our *in vivo* studies demonstrated robust tumor regression in an aggressive murine PC model treated with combined ARi/NLRP3 agonist via NLRP3-primed phagocytic TAM.

AR plays a central role in PC tumorigenesis and AR inhibition induces tumor cell apoptosis, inhibits proliferation, migration and metastasis of tumor cells^30^. Despite an initial response to ARi, PC patients ultimately demonstrate progression and develop mCRPC through various tumor cell-intrinsic mechanisms, including amplification of AR ^31,32^ and its enhancers^33,34^, AR gene point mutations^35^, AR splice variants such as AR-V7^36,37^ and the induction of AR-indifferent lineage plasticity^38^. With respect to tumor cell-extrinsic mechanisms, we have recently demonstrated that blockade of AR in CD8^+^T cells increased IFNγ expression, inhibited T cell exhaustion, restored response to anti-PD1 therapy and controlled growth of PC^39^. Our current study demonstrates that AR blockade primes NLRP3 in immunosuppressive M2 TAM, and that NLRP3 activation by small molecule NLRP3 agonist treatment controls tumor growth in an advanced murine PC model via the enhancement of phagocytosis. NLRP3 priming did not occur in low AR-expressing M1 TAM (which exhibit high *de novo* NLRP3 expression) following AR blockade therapy. However, M1 TAM demonstrated enhanced phagocytosis following NLRP3 agonist treatment, thus highlighting that low AR expression within activated anti-tumor M1 TAM could contribute to high NLRP3 expression, which can be exploited via NLRP3 agonism. Collectively, these findings demonstrate that the androgen axis suppresses NLRP3 expression and resultant innate immunity within M2 TAM.

Classically, TAM are characterized as either “antitumor” M1 or “immunosuppressive” M2. TAM have a high degree of heterogeneity and can be classified into subtypes (e.g. TREM2+ macrophages in neurodegenerative disorders, immunosuppressive IL4I1+PD-L1+IDO1+ TAM in solid tumors) based on transcriptional, metabolic and functional profiling^19,40–42^. Following AR blockade/NLRP3 agonist therapy, the NPP TAM subset phagocytosed cancer cells with corresponding upregulation of PD-L1 and CD86. Molecularly, these immune checkpoints are known to interact with PD-1/CTLA-4 on T cells, respectively which can suppress adaptive immune response, and potentially limit long-term tumor controls in cancer patients^43–45^. Taken together, these observations demonstrate that PD-L1 and CD86 are predictive biomarkers of treatment response to ADT/NLRP3 agonist combination therapy, and suggest that immuno-oncology combinations with PD-1/CTLA-4 blockade therapy could enhance adaptive anti-tumor immunity above and beyond innate immune activation.

Prior studies have demonstrated that AR signalling is a significant contributor to the growth, metastases and treatment-resistance in melanoma^46–48^, lung^49^, liver^49^, bladder^49^ and colorectal cancers^50^. Androgen-mediated sexual dimorphism is associated with superior progression-free survival in BRAF/MEKi-treated female patients and mice with melanoma, relative to males. Furthermore, AR blockade restored the anti-tumor response in male mice by tumor cell-intrinsic mechanisms^48^, suggesting a broader impact of ARi treatment on the TME to control cancer growth. In contrast, androgen-driven neutrophil maturation decreases risk of metastases in male melanoma patients and mice. This benefit was attenuated by ARi^47^, thus highlighting that sexual dimorphism is immune cell-context specific, and underscoring the importance of mechanistic studies to elucidate AR biology within different immune cell lineages. In our study, we observed low baseline NLRP3 expression within TAM of leukemia, liver, lung and pancreatic cancers in the absence of ARi treatment. Given the universality of NLRP3 priming within immunosuppressive M2 TAM following androgen blockade, this leads to the tantalizing possibility that ARi can prime NLRP3 within AR-expressing M2-TAM across multiple cancers other than PC, thereby creating a therapeutic vulnerability targetable with an NLRP3 agonist. This concept warrants further clinical investigation in neoadjuvant clinical trials across a wide range of solid malignancies.

## METHODOLOGY

### Single cell RNA sequencing and bioinformatic analysis in human samples

All human scRNAseq data were obtained from previously published repositories. These published FASTQ files were obtained from GSE143791, and further processed using CellRanger. Human genome- hg19 was considered as the reference genome to generate matrix files containing cell barcodes and transcript counts. To accurately classify cells relative to noise, unique molecular identifiers (UMI) threshold was set at 600, and these well-defined cells were included in downstream analysis. Quality control and data integration were done using Conos^51^. NLRP3 expression per cell was calculated as an average normalized (for cell size) gene expression across different cell subsets. NLRP3-high and NLRP3-low expressing populations were characterized by upper and lower quadrant of cells present in the given cell subsets, respectively. A gene set signature score was used to measure cell states in different cell subsets and conditions. Signature scores were calculated as average expression values of genes in each set. 1) Inflammasome activity, 2) M1 and 3) M2- scores in each cell were calculated using three sets of the following genes: 1) AIM2, TRAF6, TXNIP, XIAP, TNFAIP3, BCL2, CASP1, CASP12, CASP4, CARD6, CTSB, CTSV, IL18, IL1A, IL1B, IL18BP, IL33, MYD88, IRF1, IRF3, NFkB1, NFKBID, NFKBIB, NLRP3, PSTPIP1, PTGS2, PYCARD, TIRAP, TNFSF14, TNFSF4; 2) AK3, APOL1, APOL2, APOL3, APOL6, ATF3, BCL2A1, BIRC3, CCL5, CCL15, CCL19, CCL20, CCR7, CHI3L2, CSPG2, CXCL9, CXCL10, CXCL11, ECGF1, EDN1, FAS, GADD45G, HESX1, HSD11B1, HSXIAPAF1, IGFBP4, IL6, IL12B, IL15, IL2RA, IL7R, IL15RA, INDO, INHBA, IRF1, IRF7, OAS2, OASL, PBEF1, PDGFA, PFKFB3, PFKP, PLA1A, PSMA2, PSMB9, PSME2, PTX3, SLC21A15, SLC2A6, SLC31A2, SLC7A5, SPHK1, TNF, TRAIL; and 3) ADK, ALOX15, CA2, CCL13, CCL18, CCL23, CD36, CERK, CHN2, CLECSF13, CTSC, CXCR4, DCL-1, DCSIGN, DECTIN1, EGR2, FGL2, FN1, GAS7, GPR86, HEXB, HNMT, HRH1, HS3ST1, HS3ST2, IGF1, LIPA, LTA4H, MAF, MRC1, MS4A4A, MS4A6A, MSR1, P2RY14, P2RY5, SEPP1, SLC21A9, SLC38A6, SLC4A7, TGFB1, TGFBR2, TLR5, TPST2^52^. We utilized our previously published scRNAseq data of bone metastatic (BMET) PC samples (n = 7), BMET non-PC samples (n = 3) and bone marrow from patients with benign inflammation (n = 7)^16^, along with leukemia^53^ (n = 20), liver^54^ (n = 8), lung^55^ (n = 10), pancreatic^56^ (n = 21) and primary prostate cancers^57^ (n = 11) to elucidate differences in NLRP3 expression and inflammasome activity. BMET samples were obtained from ADT (gonadotropin-releasing hormone [GnRH] agonist with or without androgen receptor signaling inhibitor)-treated patients, and primary PC samples were collected from patients with no prior systemic therapies who underwent radical prostatectomy.

### Cell lines and culture conditions

iBMDM were previously generated from primary BMDM of C57BL/6 mice by transfection of J2 recombinant retrovirus carrying v-myc and v-raf oncogenes^58^ and provided by Gajewski Laboratory (University of Chicago). iBMDM cells were cultured in DMEM media containing 10% fetal bovine serum (FBS), 1% penicillin/streptomycin and 1% non-essential amino acids. Myc-CAP cells (ATCC) were cultured in DMEM media containing 10% FBS, 2% L-glutamine and 1% penicillin/streptomycin. To study the specific impact of androgens on downstream readouts of NLRP3 pathway activation and macrophage functionality, androgen enriched (AE) media was generated by adding 10% charcoal-stripped FBS (instead of 10% normal FBS) containing synthetic androgen R1881 (1 nM) to the above mentioned DMEM media. Cells were treated with AE media for 24 hours to establish androgen-regulated response.

### M1/M2-polarization

iBMDM with M0-phenotype were polarized to either M1 or M2 following 24 hours treatment with cytokine cocktails, IFNγ (5 ng/mL) + LPS (200 μg/mL) or IL-4 (20 ng/mL), respectively, and analyzed by flow cytometry for the following markers: SIRPα, CD16, MHC-II, Arg1, TIM-3, PD-L1, CD206, CD86, PD-1, CD64, and NLRP3.

### Western blot analysis

iBMDM with M0, M1, M2-phenotypes or Myc-CAP tumor cells were treated with NLRP3a and enzalutamide under AE-media conditions, as indicated in figure legends. Cells were lysed in RIPA buffer and 30 μg of total protein extracts for each sample underwent sodium dodecyl sulfate-polyacrylamide gel electrophoresis (SDS-PAGE), followed by western blot analysis. These blots were probed for NLRP3, caspase-1, cleaved caspase-1 and GAPDH.

### ELISA assay

iBMDMs were treated with NLRP3a at 1, 5, 10, 50 and 100 μM for 1 hour. Supernatants were collected and analyzed for IL-1β/IL-18 cytokine levels as per manufacturer (Thermofisher) instructions provided in ELISA kits: IL-1β (BMS6002); IL-18 (BMS618).

### Murine TME analysis

Experiments were performed in accordance with NIH guidelines and protocol approved by the Institutional Animal Care and Use Committee (IACUC) at University of Chicago. Syngeneic FVB mice that are 10-12-weeks of age (Jackson Laboratory, USA) were inoculated subcutaneously with 1x10^6^ Myc-CAP prostate cancer cells (ATCC, USA) in the right flank region. Once tumors were palpable, vernier caliper was used to monitor tumor growth.

Volumetric analysis was performed using following equation; volume (mm^3^) = 0.5 x length (mm) x width (mm) x width (mm). To identify *in vivo* NLRP3 expressing cell subsets, untreated Myc-CAP tumors were harvested at early (tumor volume = 200 mm^3^, baseline) and late (tumor volume = 2500 mm^3^, untreated) phases of tumor growth. Additionally, syngeneic mice with established Myc-CAP subcutaneous tumors (∼200 mm^3^) were treated with NLRP3a (0.3 mg/kg, *it*, single dose, NLRP3 agonist, N) for 24, 48 and 72 hours, and tumors harvested to understand dynamic changes in NLRP3 expression within TME. To assess impact of ADT on NLRP3 expression within TME, mice with established Myc-CAP subcutaneous tumors (200 mm^3^) were treated with degarelix (0.625 mg/mouse, *sc*, monthly, chemical castration, ADT), and tumors were harvested at peri- castration, early resistance, and late resistance stages (at days 7, 18 and 30, respectively).

To analyze NLRP3 expression in specific cell subsets within the TME, tumor single cell suspensions were centrifuged and incubated for 2 minutes in 1 mL of Ammonium-Chloride-Potassium (ACK) solution to lyse red blood cells. ACK was neutralized with 10 mL of PBS and the cell suspension was centrifuged at 1800 rpm for 5 minutes. The cell pellet was resuspended at density of 1x10^6^ cells/mL in PBS containing either myeloid or lymphoid antibody cocktails. Myeloid antibody cocktails were prepared by resuspending 10 μL of anti-CD45, CD11b, CD11c, Ly6c, Ly6g, MHC-II, F4/80, PD-L1, CD86 antibodies and 1 μL of Ghost-viability dye in 1 mL PBS. Lymphoid antibody cocktails were prepared by resuspending 10 μL of anti-CD45, CD3, CD4, CD8, CD19, FoxP3 antibodies and 1 μL of Ghost-viability dye in 1 mL PBS. After 30 minutes of incubation, stained single cell suspensions were washed with PBS and incubated with fix/perm buffer for 15 minutes. Following fixation, either myeloid or lymphoid antibody cocktail- stained cells were incubated overnight with 1 mL perm buffer containing 1 μg of NLRP3 antibody for intracellular stains. These cells were washed with PBS (2x) and incubated with secondary rabbit antibody (1:200 dilution in 1 mL perm buffer) for 1 hour. After completion of staining process, cells were washed, resuspended in PBS and flow cytometry was performed. Flow cytometry data were gated for myeloid and lymphoid subsets and their NLRP3 expression status assessed using FlowJo v 10.7 software.

### Bulk RNA sequencing and bioinformatic analysis

Our previously published bulk RNAseq data of mCRPC patients (n = 16)^39^ were utilized to examine correlation between AR activity and NLRP3/inflammasome activity. In this study, our mCRPC patients who progressed on enzalutamide treatment (NCT02312557, n = 3 bone sites, n = 3 liver sites, n = 6 lymph node sites and n = 4 other metastatic sites) received pembrolizumab treatment (200 mg, every three weeks, 4 cycles), and response to therapy was defined by > 25% reduction in Prostate Specific Antigen (PSA), compared to baseline. The corresponding baseline vs on-treatment FASTQ raw files were pre-processed to discard the adaptor sequences using Cutadapt (RRID:SCR_011841), and low-quality reads were eliminated. Next, filtered reads were mapped to the human genome (hg19) using STAR aligner (RRID:SCR_004463). To measure AR regulon activity of each sample, we used the VIPER R package (version 1.16.0)^59^. A log1p-transformed TPM gene expression matrix and a regulatory network (regulome) were used as inputs for single-sample VIPER analysis. The transcriptional regulatory network used in this study was curated from several databases as previously described^60^. NLRP3 expression was analyzed using log-transformed expression profile in each sample. Inflammasome activity (gene list present in the previous scRNAseq section) in each sample was determined using the single-sample gene set enrichment analysis (ssGSEA) implemented in the GSVA with R package (version 1.44.5)^61^. Similarly, inflammasome activity was also analyzed in published West Coast Dream Team (WCDT) human mCRPC mRNA dataset repository^62^.

### Repolarization assay

M1/M2-polarized iBMDM were treated with NLRP3a (10 μM, 1 hour/24 hours) and enzalutamide (10 μM, 24 hours) under AE-media conditions. Cells were trypsinized, washed with PBS (2x), stained for macrophage activation (MHC-II and CD86) and suppression (Arg1 and CD206) markers, and analyzed by flow cytometry.

### Phagocytosis assay

M1/M2-polarized iBMDM were treated with NLRP3a (10 μM, 1 hour/24 hours) and enzalutamide (10 μM, 24 hours) under AE-media. Following wash out with PBS (2x), they were co-cultured with CTV dye-stained (1:2000 dilution in PBS) Myc-CAP cells at 2:1 ratio for 1 hour. At the end of phagocytosis, cells were washed twice with PBS and stained with anti- SIRPα, CD16, MHC-II, Arg1, TIM-3, PD-L1, CD206, CD86, PD-1, CD64, and NLRP3 antibodies (1:100 dilution for each) and Ghost-viability dye (1:1000 dilution) in 1 mL PBS. Phagocytic activity of treated iBMDM was assessed by flow cytometry, and calculated by normalizing MFI of CTV dye, relative to indicated control groups in figure legends. PD-L1 and CD86 status was analyzed in phagocytic, non-phagocytic and total iBMDM.

To study potential tumor cell-intrinsic contribution of androgen blockade/NLRP3 agonist combination therapy on phagocytosis, Myc-CAP cells were treated with NLRP3a (10 μM, 24 hours) and enzalutamide (10 μM, 24 hours) under AE-media. Following treatment, cells were stained with CTV dye, and co-cultured with M1/M2-polarized macrophages at 2:1 ratio for 1 hour. Phagocytic activity of iBMDM were assessed using flow cytometry as described above.

### Cytokine array

M1/M2-polarized iBMDM were treated with NLRP3a (10 μM, 1 hour) and enzalutamide (10 μM, 24 hours) under AE-media. Supernatants were collected and analyzed using LGENDplex with macrophage 13-plexinflammatory panel.

### Murine prostate tumor growth kinetic and survival analysis

To assess anti-tumor responses, syngeneic mice with established Myc-CAP tumors (200 mm^3^) were treated with either NLRP3a (0.3 mg/kg, *it*, weekly, NLRP3 agonist, N), degarelix (0.625 μg/mouse, *sc*, monthly, chemical castration, ADT) or their combination. Tumor growth was monitored using vernier caliper for indicated timepoints in figure legends. Mice were monitored for survival from establishment of Myc-CAP tumors (∼200 mm^3^) until pre-specified cut-offs for either tumor volume (∼2500 mm^3^) or treatment duration (60 days) in both treated and untreated groups. After mice reached survival endpoint, tumors were harvested for TME analysis using flow cytometry. For acute mechanistic studies, mice with established Myc-CAP tumors were treated with above mentioned agents or their combination for 7 days. Tumors were harvested and single cell suspensions were processed for TME analysis using flow cytometry, as described above.

### *In vivo* macrophage depletion studies

To elucidate the role of macrophages in driving anti-tumor responses *in vivo*, syngeneic mice with established Myc-CAP tumors (200 mm^3^) were treated concurrently with clodronate (200 μg/mouse, *ip*, once weekly, Cl), which is known to deplete phagocytic macrophages *in vivo*^63^, singly or in combination with degarelix (0.625 μg/mouse, *sc*, monthly, ADT) plus NLRP3a (0.3 mg/kg, *it*, weekly, N) or NLRP3a monotherapy. Prostate tumor growth kinetics and TME analysis were done, as previously described.

### Proliferation assay

Myc-CAP cells were treated with NLRP3a (10 μM, 72 hours) and enzalutamide (10 μM, 72 hours) under AE-media conditions. Proliferation rate was determined using MTT assay as per manufacturer protocol.

### Data analysis

Data was analyzed using Graph Pad Prism v7. Statistical analysis was performed using one way analysis of variance (ANOVA) and Bonferroni post-test with p<0.05 level of significance. Unpaired Student t-tests were used to compare the means between two independent groups. For scRNAseq, NLRP3 expression and inflammasome activity within each cell subset were represented using Uniform Manifold Approximation and Projection (UMAP) method at single cell level. Group comparison for human data were done using Wilcoxon rank-sum tests (for two groups) or Kruskal–Wallis tests (> than two groups). Correlation analysis was done by Pearson test between two continuous variables.

### Data and code availability

The authors declare that the data/materials supporting the findings of this study are available within the Article, its Supplementary Information and public repository cited at appropriate places. Code for reproducibility/analysis of data is publicly available. All raw data are available from corresponding author upon reasonable request.

## Acknowledgements

We thank Dr. Marisa Naujokas for providing helpful suggestions for this manuscript. This work was supported by Prostate Cancer Foundation (**A.P.)** and P50CA180995 **(A.P.)**.

## Author’s contributions

**K.C.:** Conceptualization, data curation, formal analysis, validation, investigation, visualization, methodology, supervision, project administration, writing–original draft, writing-review and editing. **S.R.:** Conceptualization, data curation, formal analysis, validation, investigation, visualization, methodology, supervision, project administration, writing– original draft, writing-review and editing. **S.M.:** Data curation, formal analysis, validation, investigation, visualization, methodology, writing-review and editing. **T.H.:** Data curation, formal analysis, validation, investigation, visualization, methodology, writing-review and editing. **Y.H.:** Data curation, formal analysis, validation, investigation, visualization, methodology, writing-review and editing. **A.A.:** Data curation, formal analysis, validation, investigation, visualization, methodology, writing–original draft, writing-review and editing. **B.L.:** Data curation, formal analysis, validation, investigation, visualization, methodology, writing–original draft, writing-review and editing. **K.D.:** Writing–original draft. **S.G.:** Data curation, formal analysis, methodology. **D.K.:** Data curation, formal analysis, methodology. **R.N.:** Formal analysis, validation, methodology. **S.D.:** Formal analysis, validation, methodology. **D.Z.:** Formal analysis, validation, methodology. **Mi.L.:** Formal analysis, methodology. **F.D.:** Methodology. **R.C.:** Methodology. **J.S.:** Data curation, visualization, writing–original draft, writing-review and editing. **Ma.L.:** Methodology. **Z.X.:** Data curation, resources, visualization, methodology, supervision, writing–original draft, writing-review and editing. **D.B.S.:** Data curation, resources, visualization, methodology, supervision, writing–original draft, writing-review and editing. **A.M.:** Conceptualization, data curation, resources, investigation, visualization, methodology, supervision, writing-review and editing. **A.P.:** Conceptualization, data curation, resources, funding acquisition, investigation, visualization, methodology, supervision, project administration, writing–original draft, writing-review and editing.

