## Supplementary figure 1-11 for "Androgen blockade primes NLRP3 in macrophages to induce tumor phagocytosis"

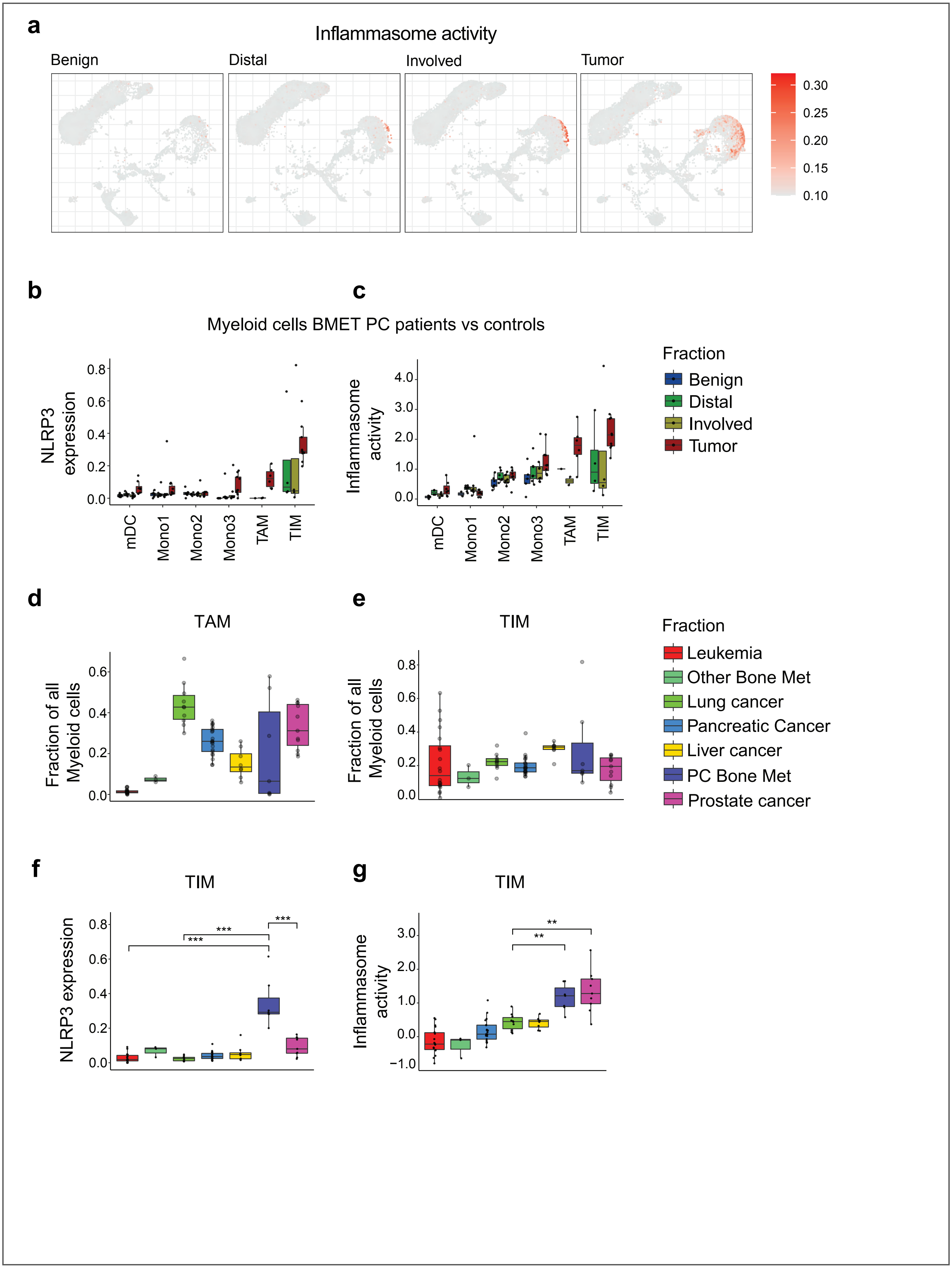
a
Inflammasome activity
Benign
Distal
Involved
Tumor
0.30
0.25
0.20
0.15
0.10
b
c
Myeloid cells BMET PC patients vs controls
NLRP3 expression
Inflammasome activity
mDC
Mono1
Mono2
Mono3
TAM
TIM
Fraction
Benign
Distal
Involved
Tumor
d
e
TAM
TIM
Fraction of all Myeloid cells
Fraction
Leukemia
Other Bone Met
Lung cancer
Pancreatic Cancer
Liver cancer
PC Bone Met
Prostate cancer
f
g
TIM
NLRP3 expression
***
Inflammasome activity
**

**Extended Data Fig. 1 │ NLRP3 is primed within TIM and TAM in ADT-treated bone metastatic PC patients, relative to other cancers and untreated primary PC. a**, scRNAseq was performed on biopsies collected from tumor, involved and distant sites of BMET PC patients as described in the Methods section, and compared with healthy benign bone marrow. Inflammasome activity was analyzed and visualized in UMAP plots. **b**-**c**, NLRP3 expression (**b**) and inflammasome activity (**c**) were quantified within myeloid subsets from tumor, involved and distant sites in BMET PC and healthy benign bone marrow. **d**-**e**, TAM (**d**) and TIM (**e**) frequencies were assessed across 7 different cancer types listed in the panels using scRNAseq and plotted. **f**-**g**, NLRP3 expression (**f**) and inflammasome activity (**g**) were plotted for tumor-inflammatory monocytes (TIM) across 7 different cancer types. Data obtained from specimens of BMET PC (n = 7 patients) for panel **a**-**g**, healthy benign bone marrow (n = 7 human donors) for panel **a**, leukemia (n = 20 patients), other bone met (non-prostate bone metastatic cancers, n = 3 patients), lung cancer (n = 10 patients), pancreatic cancer (n = 21 patients), liver cancer (n = 8 patients) and primary prostate cancer patients (n = 11 patients) for panels **d**-**g**. Except panel **a**, all panels show biological replicates as mean $\pm$ s.e. Significances/p-values were calculated by Wilcoxon rank-sum test and indicates as follows, ***$p<0.001$.

### Extended Figure 2

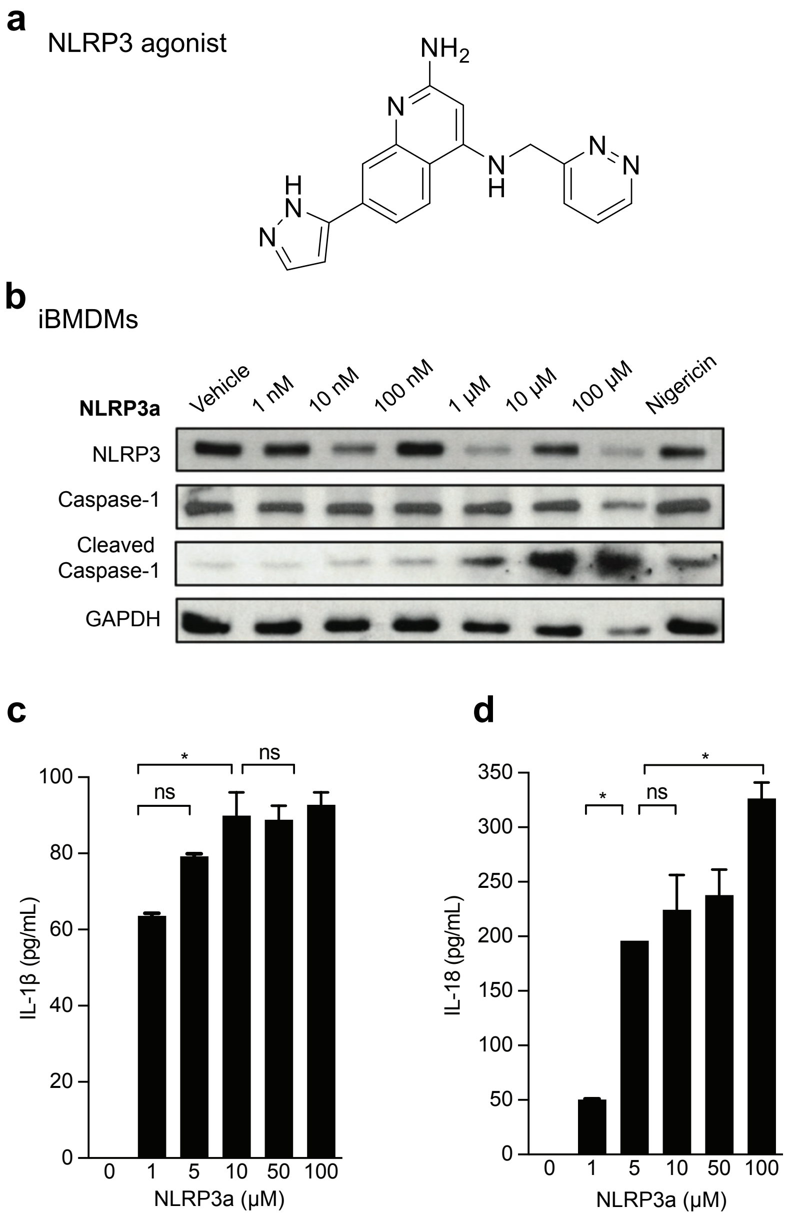

**Extended Data Fig. 2 | NLRP3a drives NLRP3 inflammasome pathway activation in immortalized bone-marrow derived macrophages (iBMDM). a**, Chemical structure of NLRP3a. **b-d**, Undifferentiated iBMDM (M0) were treated with NLRP3a in a dose-dependent/time-dependent manner as indicated, and NLRP3 inflammasome pathway activation was assessed by western blotting (**b**) and IL-1β (**c**)/IL-18 ELISA (**d**). Data obtained from n=3 biological replicates. Except panel **a** and **b**, all panels show biological replicates as mean ± s.e. Significances/p-values were calculated by one-way ANOVA and indicates as follows, *$p<0.05$; ns = not statistically significant.

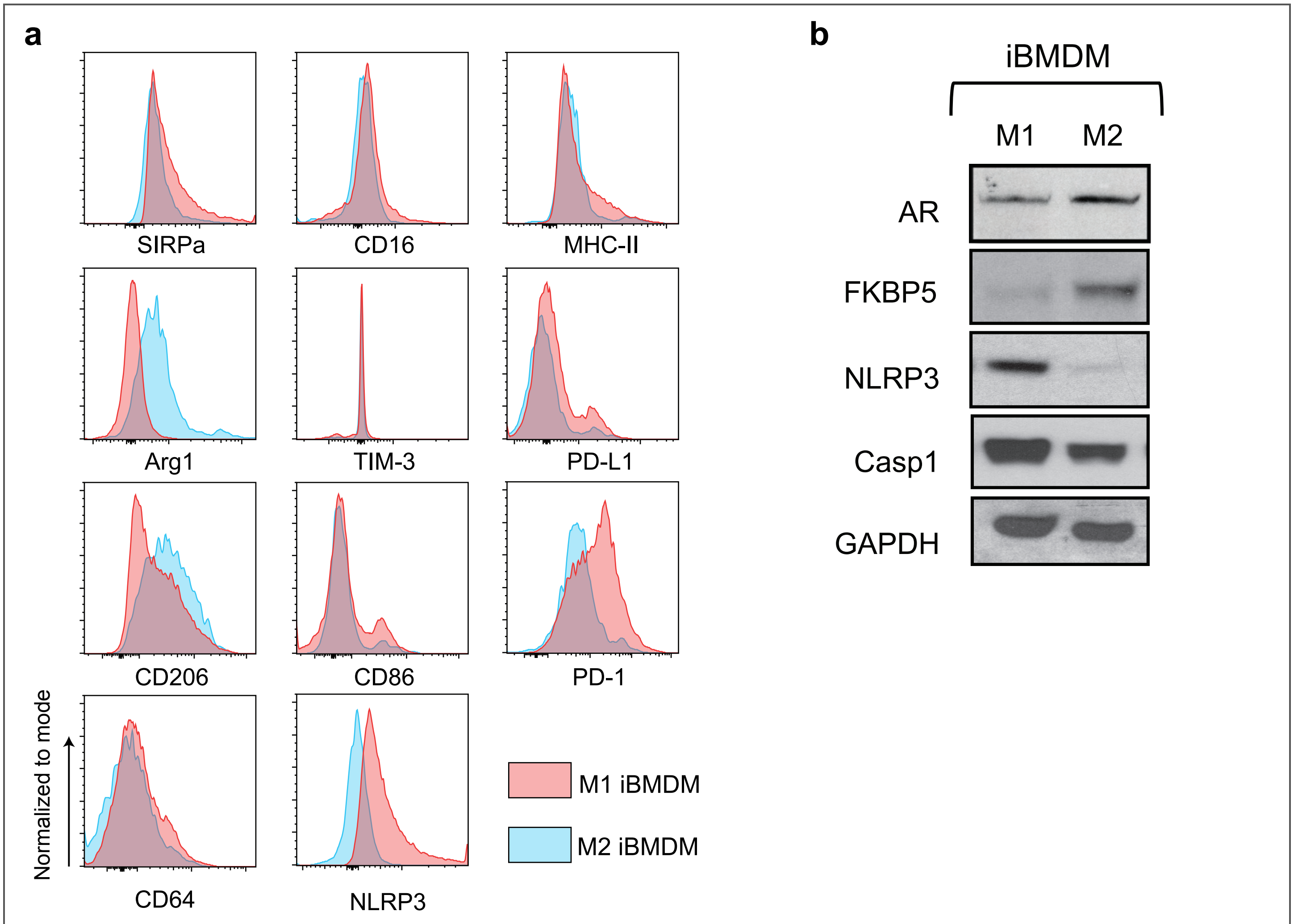
a
SIRPa
CD16
MHC-II
Arg1
TIM-3
PD-L1
CD206
CD86
PD-1
Normalized to mode
CD64
NLRP3
M1 iBMDM
M2 iBMDM
b
iBMDM
M1
M2
AR
FKBP5
NLRP3
Casp1
GAPDH

**Extended Data Fig. 3 | Anti-tumor (M1) macrophages have high NLRP3 expression with corresponding low AR expression/activity, relative to immunosuppressive (M2) macrophages. a**, iBMDM were treated with either LPS (200 ng/mL) + IFNγ (10 ng/mL) or IL-4 (20 ng/mL) for 24 hours to differentiate them into M1 or M2 phenotype, respectively, and analyzed for the indicated phenotypic markers by flow cytometry. **b**, Differential NLRP3 and AR expression/pathway activation in M1 vs M2 iBMDM was assessed by western blotting. Data obtained from the n=3 biological replicates.

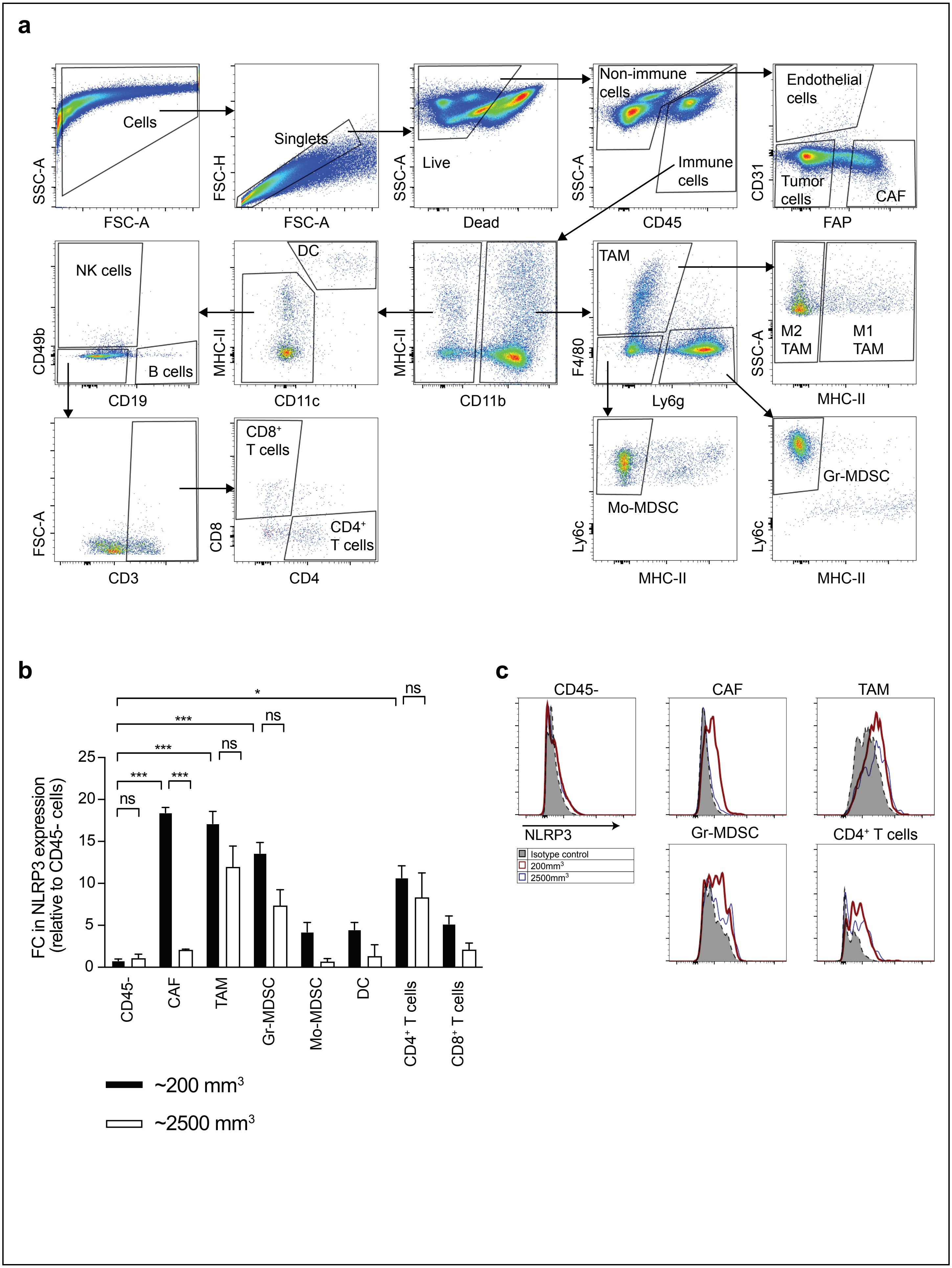
a
Cells
Singlets
Live
Non-immune cells
Immune cells
Endothelial cells
Tumor cells
CAF
SSC-A
FSC-A
FSC-H
Dead
CD45
CD31
FAP
NK cells
B cells
DC
TAM
M2 TAM
M1 TAM
CD49b
CD19
MHC-II
CD11c
CD11b
F4/80
Ly6g
SSC-A
CD8+ T cells
CD4+ T cells
Mo-MDSC
Gr-MDSC
FSC-A
CD3
CD8
CD4
Ly6c
b
FC in NLRP3 expression (relative to CD45- cells)
ns
***
*
0
5
10
15
20
25
CD45-
CAF
TAM
Gr-MDSC
Mo-MDSC
DC
CD4+ T cells
CD8+ T cells
~200 mm3
~2500 mm3
c
CD45-
CAF
TAM
Gr-MDSC
CD4+ T cells
NLRP3
Isotype control
200mm3
2500mm3

**Extended Data Fig. 4 | NLRP3 is expressed in immune cells and cancer-associated fibroblasts within c-Myc driven murine prostate tumor microenvironment, but not in cancer cells. a-c**, Syngeneic FVB mice with established Myc-CAP tumors were harvested at early (~200 $mm^3$) and late phases (~2500 $mm^3$) of tumor growth. Tumor immune vs non-immune populations were analyzed by flow cytometry and representative gating strategies are depicted (**a**). Bar graphs show NLRP3 expressing immune populations, relative to tumor cells ($CD45^-$ cells) within TME (**b**). Histograms demonstrate NLRP3 expression in immune vs non-immune populations at early-/late-stages of Myc-CAP tumors, relative to isotype controls (**c**). Data obtained from n = 3-5 mice per group. Panel **b** shows biological replicates as mean $\pm$ s.e. Significances/p-values were calculated by Student t-test and indicates as follows, *$p<0.05$ and ***$p<0.001$; ns = not statistically significant. FC = fold change.

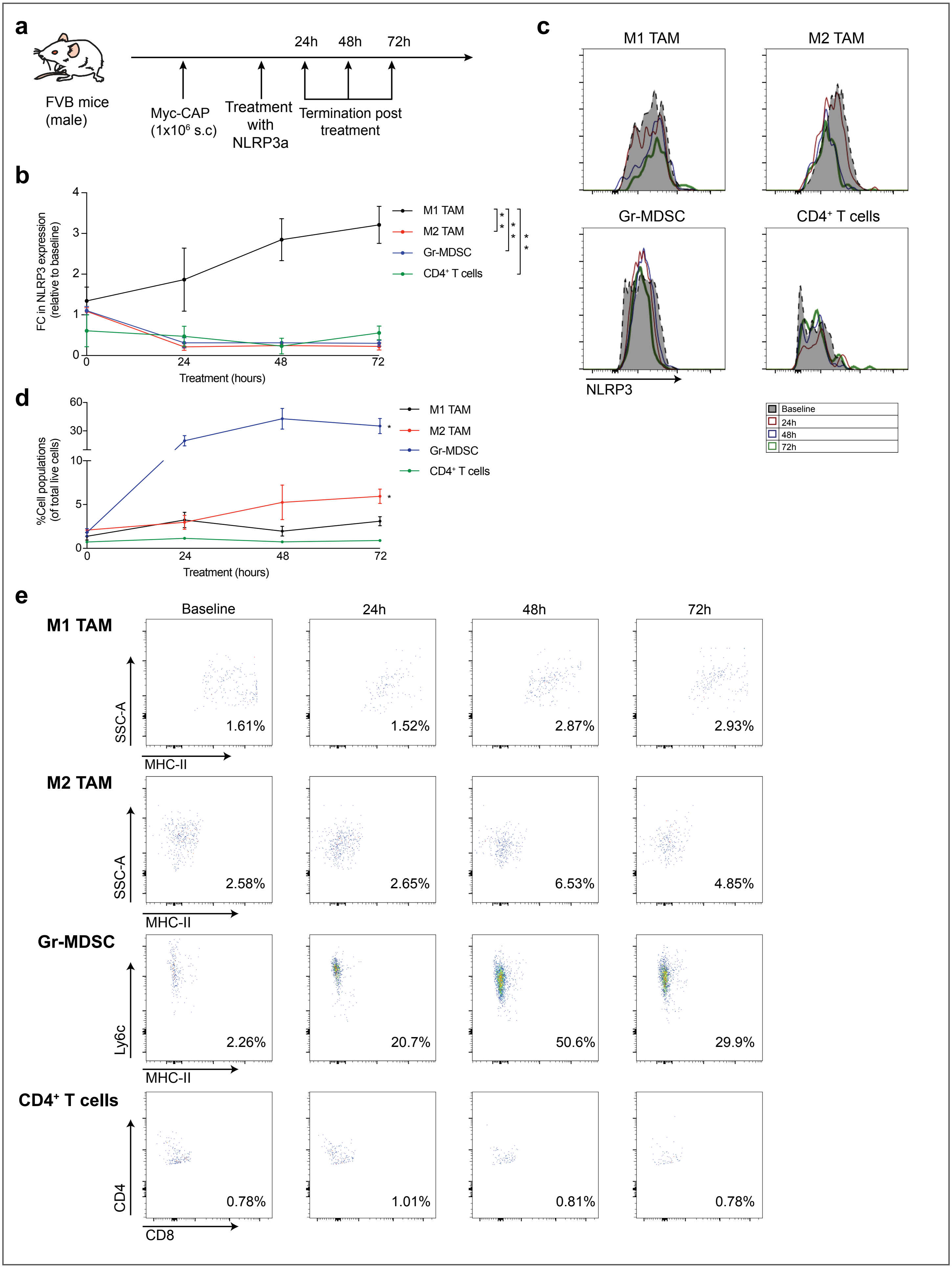
a
FVB mice (male)
24h 48h 72h
Myc-CAP (1x10⁶ s.c)
Treatment with NLRP3a
Termination post treatment
b
FC in NLRP3 expression (relative to baseline)
Treatment (hours)
M1 TAM
M2 TAM
Gr-MDSC
CD4⁺ T cells
c
M1 TAM
M2 TAM
Gr-MDSC
CD4⁺ T cells
NLRP3
Baseline
24h
48h
72h
d
%Cell populations (of total live cells)
Treatment (hours)
M1 TAM
M2 TAM
Gr-MDSC
CD4⁺ T cells
e
Baseline
24h
48h
72h
M1 TAM
SSC-A
MHC-II
1.61%
1.52%
2.87%
2.93%
M2 TAM
SSC-A
MHC-II
2.58%
2.65%
6.53%
4.85%
Gr-MDSC
Ly6c
MHC-II
2.26%
20.7%
50.6%
29.9%
CD4⁺ T cells
CD4
CD8
0.78%
1.01%
0.81%
0.78%

**Extended Data Fig. 5 | M1 TAM have high NLRP3 expression following acute NLRP3 agonist treatment, relative to other immune cell subsets. a**, Schema illustrating that syngeneic Myc-CAP tumor-bearing mice (~200 $mm^3$, baseline) were treated with NLRP3a (0.3 mg/kg, *it*) for the indicated time-points (24/48/72h). **b**-**e**, tumors were harvested and analyzed for NLRP3 expression (**b**) in M1/M2 TAM, Gr-MDSC and $CD4^+$T cells by flow cytometry. Representative histograms show NLRP3 expression in these cells (**c**). Furthermore, frequency of these cells (relative to total live cells), were analyzed within the TME following NLRP3a treatment by flow cytometry (**d**) and representative flow plots are indicated (**e**). Data obtained from n = 3-5 mice per group. Panel **b** and **d** show biological replicates as mean $\pm$ s.e. Significances/p-values were calculated by Student T-test. In panel B, a statistical comparison was done for NLRP3 expression in M1-TAM relative to other immune cell populations at 72 hours following NLRP3a treatment. In panel **d**, a statistical comparison was done for changes in cell frequency at 72 hours relative to 0 hour following NLRP3a treatment and indicates as follows, *p<0.05 and **p<0.01; ns = not statistically significant. FC = fold change.

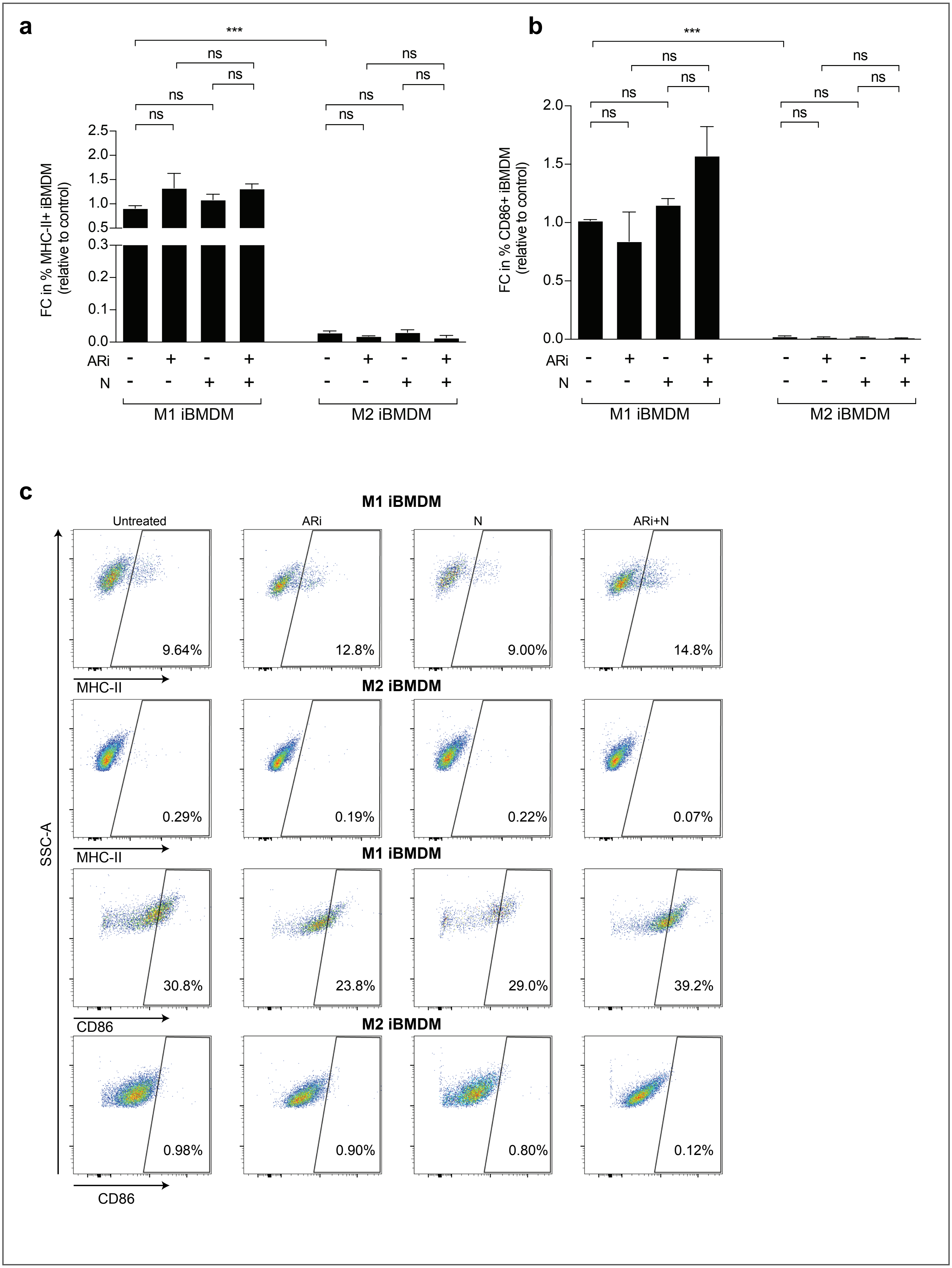
a
b
c
***
ns
FC in % MHC-II+ iBMDM (relative to control)
FC in % CD86+ iBMDM (relative to control)
ARi
N
M1 iBMDM
M2 iBMDM
Untreated
ARi
N
ARi+N
9.64%
12.8%
9.00%
14.8%
0.29%
0.19%
0.22%
0.07%
30.8%
23.8%
29.0%
39.2%
0.98%
0.90%
0.80%
0.12%
MHC-II
CD86
SSC-A

**Extended Data Fig. 6 | ARi/NLRP3a treatment does not phenotypically polarize M2 iBMDM into M1 iBMDM. a-c**, iBMDM were differentiated into M1 and M2 phenotypes using cytokine cocktail, as described in the Methods section, and treated with enzalutamide (ARi, 10μM, 24 hours), NLRP3a (10μM, 1 hour) or their combination. Flow cytometry was performed to quantify MHC-II (**a**) and CD86 (**b**) expressing iBMDM. Representative flow cytometry plots depict quantification of activated iBMDM, defined as MHC-II or CD86 expressing iBMDM relative to total live M1/M2 iBMDM (**c**). Data obtained from n = 3 biological replicates. Panels **a** and **b** show biological replicates as mean $\pm$ s.e. Significances/p-values were calculated by one-way ANOVA and indicates as follows, ns = not statistically significant. FC = fold change. Control = untreated M1 iBMDM group.

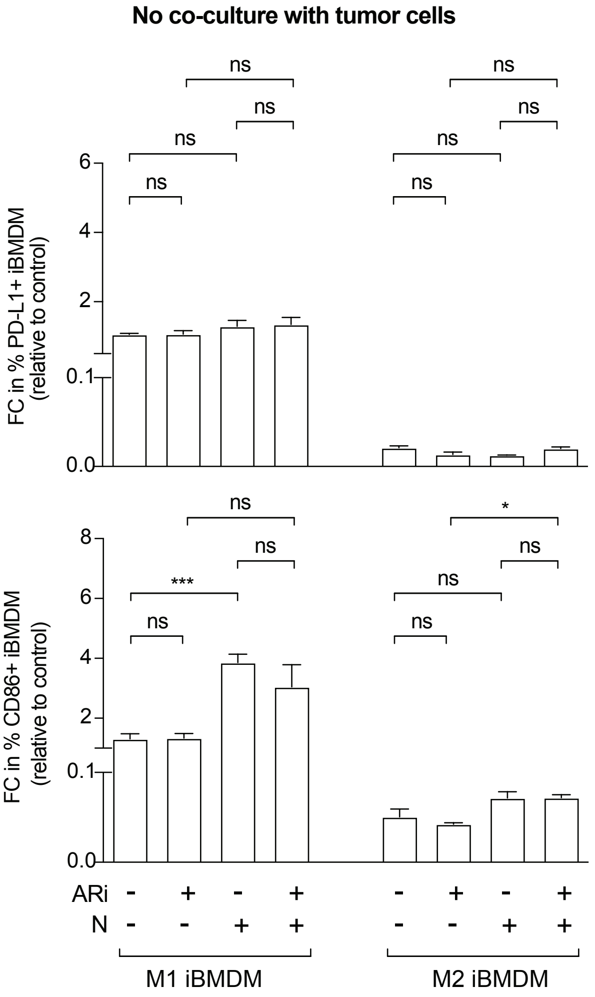
No co-culture with tumor cells
FC in % PD-L1+ iBMDM (relative to control)
FC in % CD86+ iBMDM (relative to control)
ns
***
*
ARi
N
M1 iBMDM
M2 iBMDM

**Extended Data Fig. 7 | M1/M2 iBMDM do not express PD-L1 and CD86 following *in vitro* AR blockade/NLRP3a combination, in the absence of co-culture with tumor cells.** iBMDM were differentiated into M1 or M2 phenotypes using cytokine cocktail, as described in the Methods section, and treated with enzalutamide (ARi, 10μM, 24 hours), NLRP3a (10μM, 24 hours) or their combination. M1/M2 iBMDM were profiled using flow cytometry to quantify PD-L1 and CD86 expressing populations. Data obtained from n = 3 biological replicates. All panels show biological replicates as mean $\pm$ s.e. Significances/p-values were calculated by one-way ANOVA and indicates as follows, *p<0.05 and ***p<0.001; ns = not statistically significant. FC = fold change. Control = untreated M1 iBMDM group.

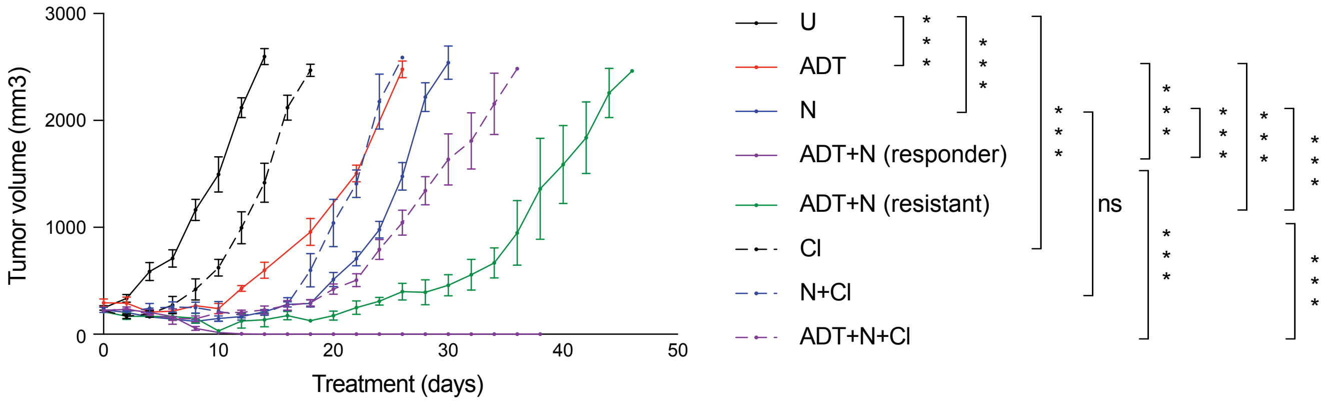
3000
2000
1000
0
Tumor volume (mm3)
0
10
20
30
40
50
Treatment (days)
U
ADT
N
ADT+N (responder)
ADT+N (resistant)
CI
N+CI
ADT+N+CI
***
ns

**Extended Data Fig. 8 | AR blockade/NLRP3a drives tumor clearance in syngeneic Myc-CAP tumor-bearing mice via a tumor cell non-autonomous mechanism.** Syngeneic FVB mice with established Myc-CAP tumors (~200 $mm^3$) were treated with degarelix (ADT, 0.625µg, *sc*), NLRP3a (0.3 mg/kg, *it*) or their combination either until non-palpable tumors for 4 weeks or until tumors reached euthanasia end-point (~2500 $mm^3$). In parallel, corresponding cohorts of mice received clodronate (200µg, *ip*, 1-hour pretreatment), for systemic depletion of phagocytic TAM, prior to administration of N and ADT/N. Tumor volumes were measured using Vernier caliper and plotted. Data obtained from n = 6-9 mice per group. Figure show biological replicates as mean ± s.e. Significances/p-values were calculated by two-way ANOVA and indicates as follows, ***p<0.001 and ns = not statistically significant. U = untreated tumors.

**a**

| Macrophages | Cytokines |
|---|---|
| M1 macrophages | IFNα, IFNβ, IFNγ, TNFα, GM-CSF, CXCL1, CXCL10, CCL2, CCL5, IL-1β, IL-6, IL-10, IL-12p70 |
| M2 macrophages | CCL2, IL-10 |

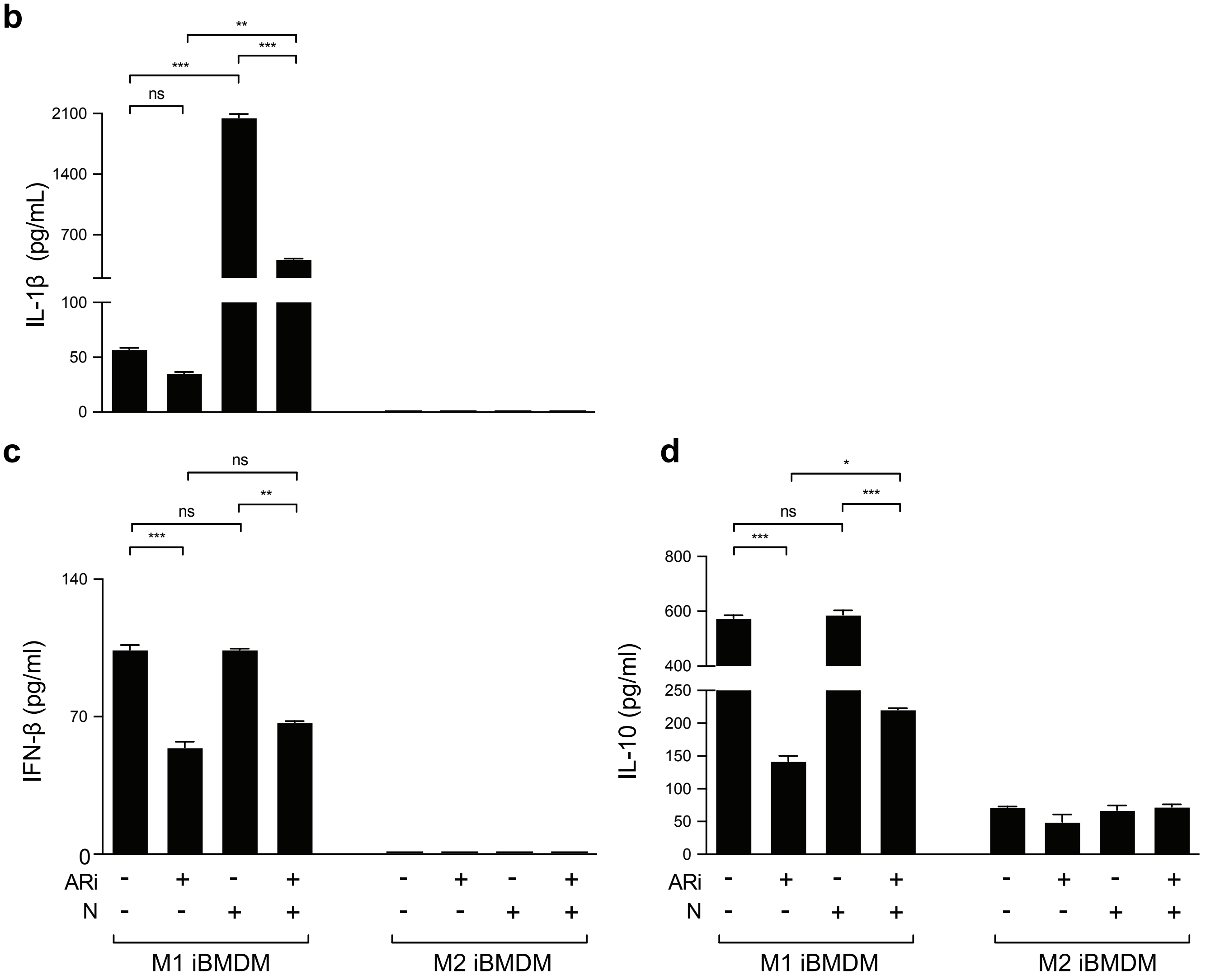

**Extended Data Fig. 9 | AR blockade/NLRP3 agonist combination does not result in pro-inflammatory cytokine release from M2 iBMDM. a-d**, iBMDM were differentiated into M1 and M2 phenotypes and treated with enzalutamide (ARi, 10μM, 24 hours), NLRP3a (10μM, 1 hour) or their combination. Cytokine array was performed on supernatants collected from the indicated treatment groups and analyzed for changes in the inflammatory secretome. M1 or M2 iBMDM secreted cytokines that were measured are listed for reference (**a**). IL-1β (**b**), IFN-β (**c**) and IL-10 (**d**) were plotted to demonstrate impact of AR blockade/NLRP3 agonist treatment on cytokine release. Data obtained from n = 3 biological replicates for *in vitro* studies. Except panel **a**, all panels show biological replicates as mean ± s.e. Significances/p-values were calculated by one-way ANOVA and indicates as follows, *p<0.05, **p<0.01 and ***p<0.001; ns = not statistically significant. Detection limit in the assays are 1.54, 1.54 and 15.1 for IL-1β, IFN-β and IL-10 cytokines, respectively.

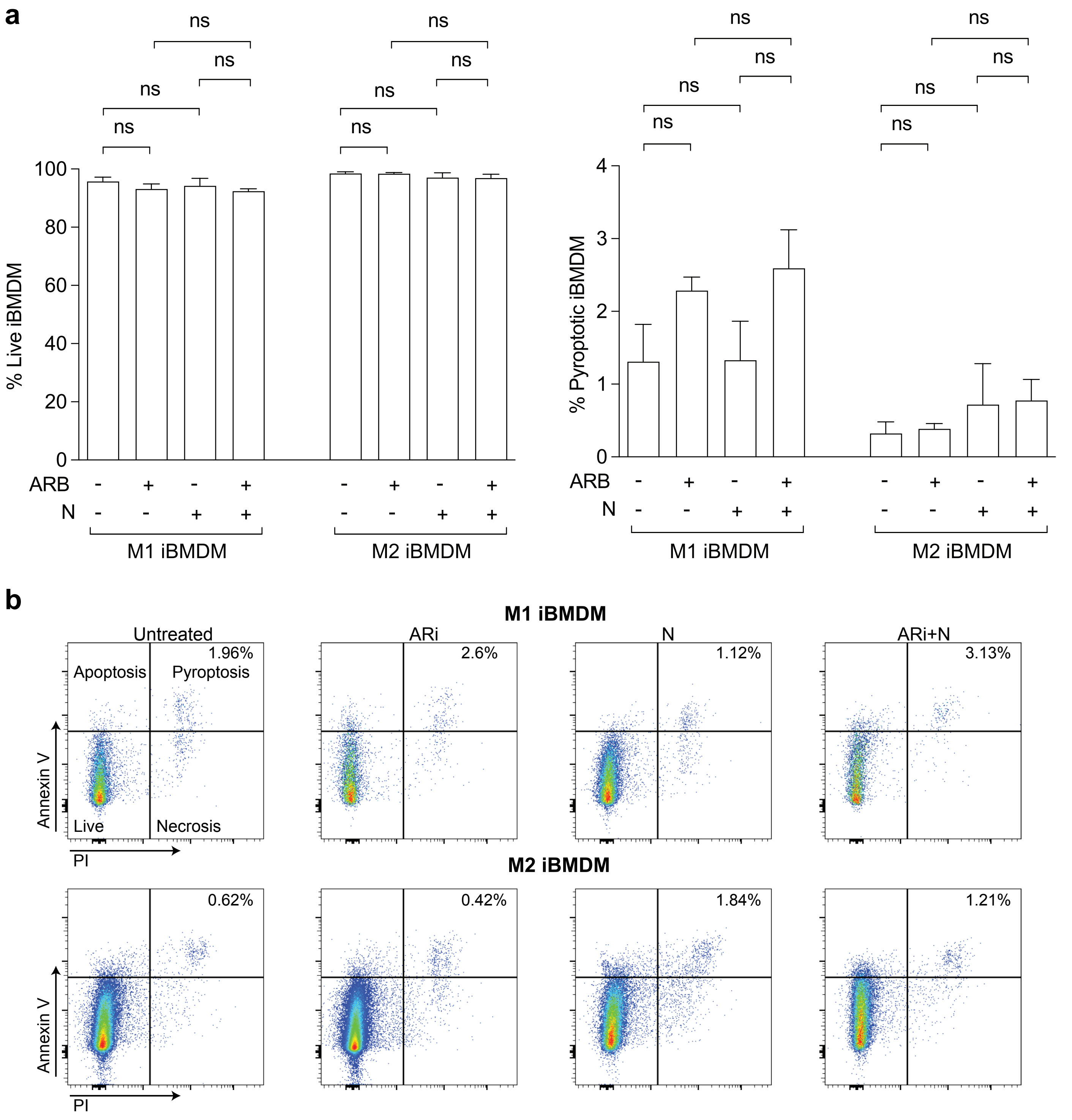
a
ns
ns
ns
ns
% Live iBMDM
100
80
60
40
20
0
ARB
N
M1 iBMDM
M2 iBMDM
% Pyroptotic iBMDM
4
3
2
1
0
b
M1 iBMDM
Untreated
ARi
N
ARi+N
1.96%
2.6%
1.12%
3.13%
Apoptosis
Pyroptosis
Live
Necrosis
Annexin V
PI
M2 iBMDM
0.62%
0.42%
1.84%
1.21%

**Extended Data Fig. 10 | ARi/NLRP3a combination does not induce pyroptotic cell death in M1/M2 iBMDM. a-b**, iBMDM were differentiated into M1 and M2 phenotypes using cytokine cocktail, as described in the Methods section, and treated with enzalutamide (ARi, 10μM, 24 hours), NLRP3a (10μM, 24 hours) or their combination. Flow cytometry was performed to quantify viable (AnnexinV$^-$/PI$^-$) and pyroptotic (AnnexinV$^+$/PI$^+$) M1/M2 iBMDM (**a**). Representative flow cytometry plots depicting pyroptotic status of M1/M2 iBMDM (relative to total single M1/M2 iBMDM) following ARi+/-NLRP3a treatment vs untreated control (**b**). Data obtained from n = 4 biological replicates. Panel **a** shows biological replicates as mean $\pm$ s.e. Significances/p-values were calculated by one-way ANOVA and indicates as follows, ns = not statistically significant.

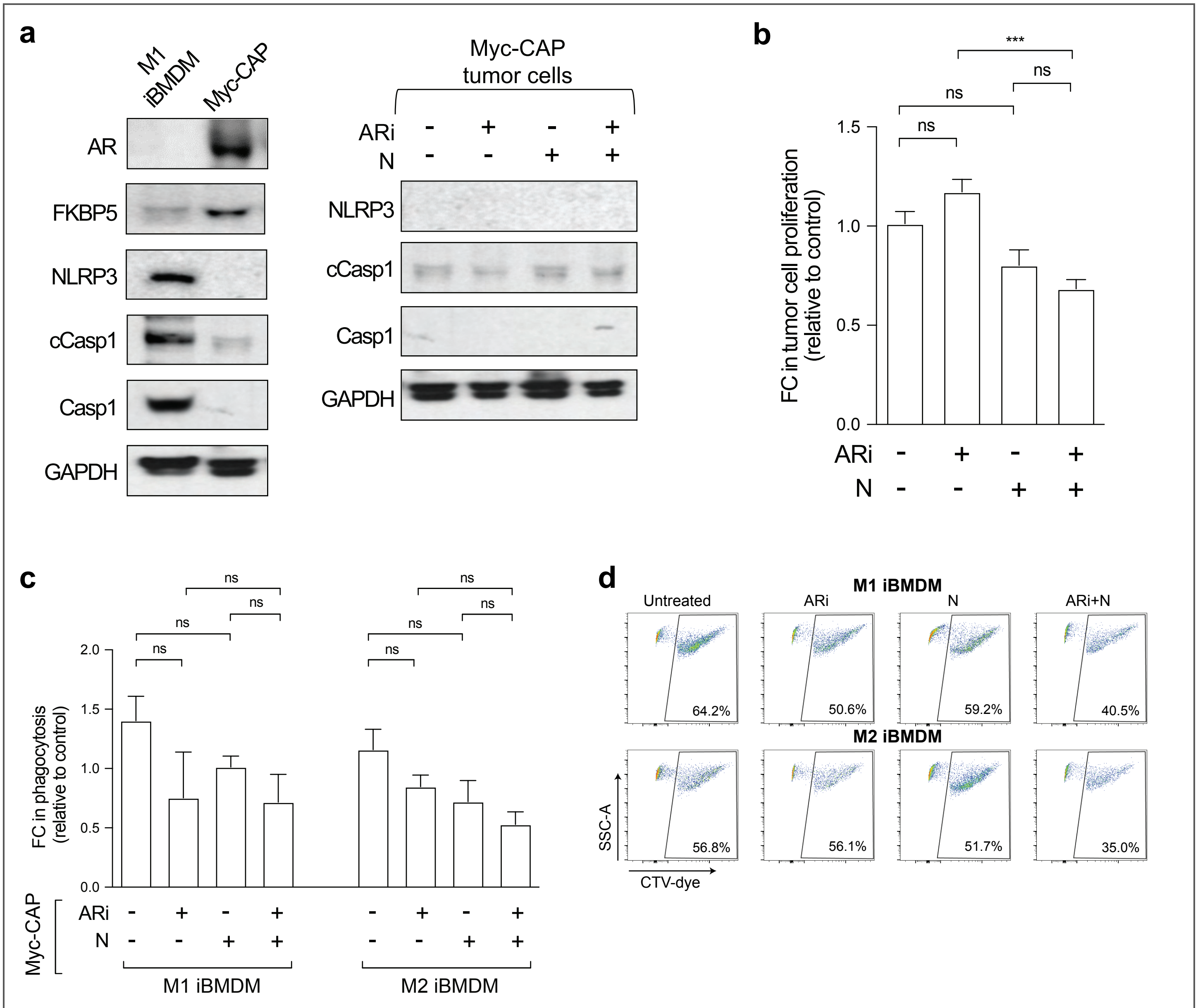
a
M1 iBMDM
Myc-CAP
AR
FKBP5
NLRP3
cCasp1
Casp1
GAPDH
Myc-CAP tumor cells
ARi - + - +
N - - + +
NLRP3
cCasp1
Casp1
GAPDH
b
FC in tumor cell proliferation (relative to control)
***
ns
ns
ns
1.5
1.0
0.5
0.0
ARi - + - +
N - - + +
c
FC in phagocytosis (relative to control)
ns
ns
ns
ns
2.0
1.5
1.0
0.5
0.0
Myc-CAP
ARi - + - + - + - +
N - - + + - - + +
M1 iBMDM
M2 iBMDM
d
M1 iBMDM
Untreated
ARi
N
ARi+N
64.2%
50.6%
59.2%
40.5%
M2 iBMDM
56.8%
56.1%
51.7%
35.0%
SSC-A
CTV-dye

**Extended Data Fig. 11 │ AR blockade/NLRP3a drives tumor clearance in syngeneic Myc-CAP tumor-bearing mice via a tumor cell non-autonomous mechanism. a**, Myc-CAP cells were treated with enzalutamide (ARi, 10 μM, 24 hours), NLRP3a (10 μM, 1 hour) or their combination. Western blotting was performed to probe for AR and NLRP3 inflammasome pathway activation. **b**, Myc-CAP cells were treated with ARi (10 μM), N (10 μM) or their combination for 72 hours, and their proliferation rates were determined using MTT assay. **c-d**, iBMDM were differentiated into M1 and M2 phenotypes and co-cultured with enzalutamide (10 μM, 24 hours), NLRP3a (10 μM, 24 hours), or enzalutamide/NLRP3a-treated CTV-dye stained Myc-CAP cells for 1 hour. Flow cytometry was performed to count CTV-dye$^{+}$iBMDM, indicative of phagocytosis, and represented as fold change relative to control (**c**). Representative flow cytometry plots depicting phagocytic status of M1/M2 iBMDM (relative to total live M1/M2 iBMDM, (**d**)), gated as shown in Fig. 4**b**. Data obtained from n = 3 biological replicates. Except for panel **a**, **d**, all panels show biological replicates as mean ± s.e. Significances/p-values were calculated by one-way ANOVA (panel **b**, **c**) and indicates as follows, ***p<0.001 and ns = not statistically significant. FC = fold change. Control = untreated tumor cells and M1 iBMDM groups for panel **b** and **c**, respectively.
